# Dual-Mode Microfluidic Immunostaining (Dumi) Device for Diagnostic Biomarkers Detection and Tumor Microenvironment Evaluation

**DOI:** 10.1101/2025.07.14.664679

**Authors:** Yu Zhang, Yanhua Huang, Jing Jia, Xiaoguang Guo, Fuxiu Liu, Bing Shi, Zhen-li Huang

## Abstract

The limited availability of tissue samples from rare tumors poses a major barrier to advances in precise diagnosis, prognostic evaluation, and therapeutic research—challenges exemplified by primary central nervous system diffuse large B-cell lymphoma (PCNS-DLBCL). However, both immunohistochemical (IHC) subtyping and preliminary studies of the tumor microenvironment (TME) using multiplex immunolabeling rely on conventional staining workflows that consume numerous serial sections per case, leading to irreversible depletion of precious samples from rare tumors. Here, we developed a dual-mode microfluidic immunostaining (Dumi) device, which integrates diagnostic and research workflows into a single, automated platform. Clinical validation shows that > 90% reduction in section usage, complete detection of subtyping diagnostic biomarkers on just 1–2 slides, and displays the TME maps incorporating six immune markers for analysis research. This microfluidic innovative offers an efficient, cost-effective, and multifunctional alternative immunostaining methods, overcoming the limitations of scarce tissue resources and providing a clinically accessible solution to the diagnostic and therapeutic challenges with rare tumors.

**Teaser:** Dumi device enables multiplex IHC & TME mapping on 1–2 slides, reducing tissue use by >90% for rare tumors like PCNS-DLBCL.

## Introduction

The precise clinical diagnosis and treatment of rare tumors have long been constrained by the fundamental contradiction between the limited availability of tissue samples and the need for multidimensional analysis(*1, 2*). This contradiction is particularly evident in the study of primary central nervous system diffuse large B-cell lymphoma (PCNS-DLBCL) (*3-5*). As a rare and high aggressiveness subtype of diffuse large B-cell lymphoma (DLBCL) characterized by complex diagnostic requirements (*6, 7*). Pathological subtyping of PCNS-DLBCL necessitates immunohistochemical (IHC) staining for dozens of biomarkers. However, stereotactic biopsy typically yields only a small amount of tissue, and conventional IHC on a single section can report only one biomarker, putting conventional diagnostic workflows at the risk of tissue depletion (*8, 9*). While automated staining systems have accelerated the efficiency of batch processing, optimized workflow management, and saved labor and time costs; neither manual methods nor automated stainers can fundamentally overcome the challenge of limited tissue availability.

More importantly, the diagnosis-first model of resource allocation compels scientific research, such as studies on the tumor microenvironment (TME) of rare tumors, to a secondary priority. In rare tumors, in-depth analysis of immune cell infiltration patterns and key regulatory molecular networks within the TME can yield valuable multidimensional data on the spatial distribution biomarkers, supporting lymphoma subtyping and prognostic evaluation (*10, 11*). Among patients with PCNS-DLBCL, approximately 35% to 60% experience treatment failure after first-line therapy, including primary non-response and relapse (*12, 13*). This treatment failure may be linked to an immunosuppressive microenvironment and chemotherapy resistance driven by metabolic abnormalities of tumor cells. However, such findings have yet to be translated into clinically applicable synchronous diagnostic tools, primarily because conventional techniques cannot simultaneously perform multidimensional analysis on limited tissue samples. In conventional clinical practice, diagnostic and research workflows operate independently, with large amounts of tissue consumed during initial pathological subtyping. Although single-slide multiplex immunofluorescence (mIF) for TME profiling has been demonstrated, it still relies on multiple preliminary single marker validations, leading to further sample depletion and extended detection times. Stepwise detection also introduces significant data heterogeneity (*14*). Furthermore, existing multiplexed antibody-based imaging technologies entail high costs related to imaging platforms, reagent consumption, and specialized personnel. They also involve complex, labor-intensive procedures and often rely on proprietary dyes and labeling techniques, making reproducibility and standardization across different laboratories challenging. These limitations amplify the risks of working with rare tissue samples (*15*). Sample scarcity constrains TME analysis, and the consequent lack of TME data severely hinders the optimization of personalized treatment strategies. This contributes to a limited understanding of rare cases and exacerbates clinical diagnostic difficulties. Therefore, conserving rare tissue samples for subsequent scientific research remains a pressing and unresolved challenge.

Microfluidic technology has brought revolutionary breakthroughs to pathological analysis by enabling high-sensitivity biomarker detection and achieving single-cell resolution. For example, on-chip multiplex immunohistochemistry (mIHC) systems that utilize multiple tissue regions can sequentially stain over ten biomarkers on a single section, significantly reducing time and resource consumption while minimizing manual labor and variability compared to conventional approaches (*16*). However, microfluidic devices applied to tissue sections remain limited to a single detection modality. One type is used for multi-region tissue multiplexing to enable automated biomarkers detection. For example, Aditya Kashyap et al. proposed a microfluidic probe (MFP) capable of performing flexible and rapid IHC staining on submillimeter regions of tissue for high-precision quantification and grading assessment of breast cancer markers (*17*). Chang Hyun Cho et al. introduced the “barcodes-IHC” concept based on multiplex microchannels, in which linear patterns of biomarkers barcodes to enable quantitative analysis of four breast cancer biomarkers (*18*). The other is used for whole-slide mIF to map the TME. Daniel Migliozzi et al. proposed a microfluidic method that integrates multiple cycles of antibody staining and fluorescence signal elution on a single slide, achieving the identification of ten immuno-markers in pancreatic cancer (*19*). Andrea J. Radtke et al. advanced the iterative bleaching extends multiplexity (IBEX) technology by developing an automated microfluidic staining platform. Their method used borohydride derivates to bleach fluorescently conjugated antibodies and enhancing tissue adhesion to achieve 16-plex labeling in human kidney tissue (*20*). However, existing studies have not yet achieved a direct integration between the initial diagnostic marker screening and TME reconstruction. The disconnect between diagnostic efficiency and TME analysis reveals a critical gap in current technology: an integrated platform designed to extract multidimensional data from limited tissue samples, with full compatibility for clinical workflows.

In this study, we developed a dual-mode microfluidic immunostaining device (Dumi) for histopathological sections, which, for the first time, integrates rapid mIHC diagnostic screening and high-resolution TME analysis based on mIF images on just 1-2 tissue sections. By innovatively combining a 3D-printed manifold with a two-layer microfluidic chip, we engineered a three-dimensional microfluidic device featuring open-loading reagent reservoirs and sequential reagent interaction, and highly integrated immunostaining module capable of flexibly handling multiple fluid streams. This configuration enables a continuous, automated workflow that seamlessly switches between two functional modes: (1) Rapid multi-channel immunostaining, utilizing multiple tissue regions to perform fast, sequential mIHC staining for diagnostic biomarkers detection; (2) High-resolution whole-chamber immunostaining, where multi-channel staining results guide the selection of biomarkers for whole-slide mIF staining, enabling the spatial localization of in situ microenvironmental cell communities. We characterized the performance of mIHC and mIF staining in both modes, and optimized flow rates and times to establish an efficient Dumi-based workflow. Considering the diffuse cellular infiltration characteristic of PCNS-DLBCL, we initially applied Dumi to DLBCL to minimize heterogeneity-induced bias in single-channel assays, and demonstrated its scalability in tissues exhibiting comparatively low diagnostic marker heterogeneity. Clinical validation demonstrated that, compared to conventional immunostaining methods, Dumi reduces the tissue section consumption for DLBCL diagnosis and mIF experiments from 20-30 sections down to 1-2, while still providing analytically valuable rapid IHC diagnostic results. Furthermore, a single section of PCNS-DLBCL processed within the Dumi, we reconstructed the TME map incorporating six biomarkers and analyzed the relative spatial distribution characteristics among different cell populations. Dumi combines minimal resource consumption with a user-friendly design, offering a low-cost and standardized tool for resource-limited settings, seamlessly integrating multidimensional data quantification into existing diagnostic workflows. It provides an automated, resource-saving solution for personalized precision medicine in rare tumors, accelerating the discovery of novel therapeutic strategies and diagnostic tools, while improving patient outcomes and reducing healthcare costs. Importantly, Dumi exhibits excellent reagent compatibility and multi-platform versatility, which positively facilitates the clinical translation and application of multidimensional TME spatial analysis.

## Results

### Overview

To enable simultaneous detection of multiple diagnostic biomarkers and TME analysis on a single section, we developed and fabricated Dumi, a platform capable of multi-channel parallel immunostaining for 16 biomarkers and multiple cycles of iterative whole-chamber immunostaining. Dumi is directly compatible with clinically available FFPE residual samples and conventional microscopy platforms, and follows the same preprocessing protocols as standard IHC diagnostics (Fig. 1A). The microfluidic chip includes a 3D-printed fluidic exchange manifold with internal channels (Fig. 1B(i) and Fig. S1), featuring 16 open-loading reservoirs for customizable reagents loading. The yellow-highlighted channels serve as central reagent feeds for automated injection and flexible distribution of reagents via a switching valve. The lower part of the 3D-printed manifold contains a recessed area designed to accommodate the two-layer tissue staining microfluidic chip and slide, with the chip consisting of an air valve layer and a biomarker staining layer (Fig. 1B(ii)). The biomarker staining layer, comprised of 16 equal-length microchannels (300 μm wide and 80 μm high), directly contacts the tissue section. The air valve layer operates in both positive and negative pressure modes, allowing dynamic switching between 16 isolated microchannels or entire chamber, thereby enabling control of both multi-channel immunostaining and whole-chamber immunostaining modes (Fig. 1B(iii)). The chip is reversibly sealed via a pressure-based clamping mechanism and can be mounted onto a microscope stage, enabling Dumi to be transferred seamlessly between imaging platforms for real-time observation (Fig. S2 and Fig. S3). Two immunostaining strategies were implemented: brightfield mIHC and tyramide signal amplification (TSA)-based mIF (*21-23*). The microfluidic staining advantage arises because, in conventional static incubation, antibody concentration near the solid-liquid interface decreases, forming a depletion zone, while antibodies outside this zone diffuse freely, resulting in inefficient replenishment. In contrast, the advection in the microfluidic system confines molecular movement primarily to the flow direction, facilitating efficient delivery of antibodies to the solid surface and replenishment of the depletion zone. This significantly enhances antigen–antibody binding efficiency (Fig. 1B(iv)), reducing diffusion distances and improving reaction kinetics so that incubation times of over 1 h are shortened to under 10 min, and also improved the overall time of the process (Table S1). Moreover, continuous flow rapidly removes unbound antibodies, further improving staining uniformity (*24*).

**Fig. 1.**
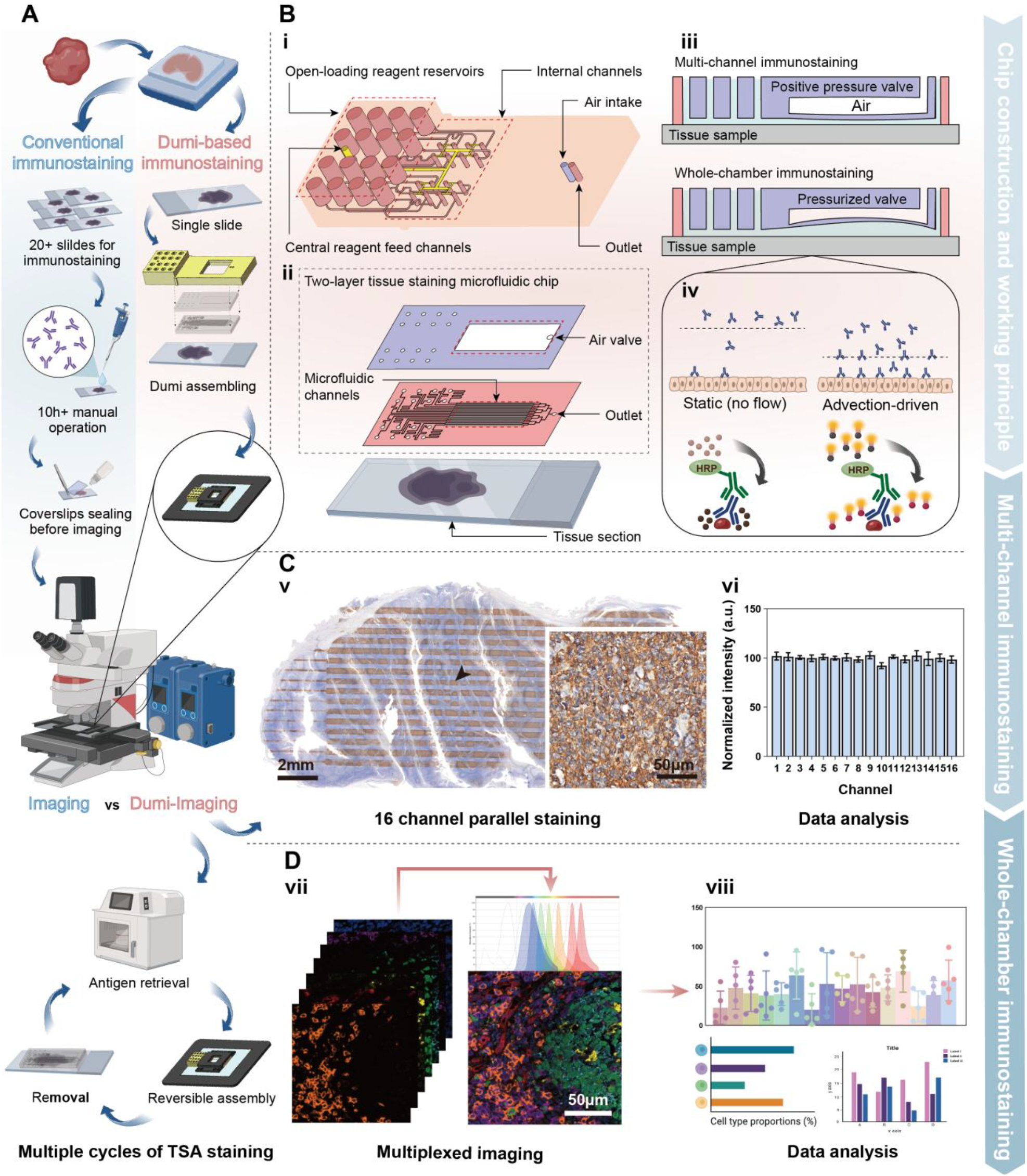
Structure, working principle, operational modes, and workflow of Dumi (created with BioRender.com). **(A)**Comparison of workflows: conventional immunostaining versus Dumi-based immunostaining. **(B)**Structure and working principle of Dumi: (i) 3D-printed fluidic exchange manifold with 16 open-loading reagent reservoirs for individual antibody solutions. These reservoirs are internally connected to the inlets of the two-layer tissue staining microfluidic chip. A central reagent feed channels have one inlet port for introducing additional reagents. The reagent feed inlet port is connected to all 16 internal channels via branching distribution channels. (ii) Two-layer tissue staining microfluidic chip. The upper air valve layer and the lower biomarker staining layer containing 16 parallel microchannels above the tissue section. (iii) Operational modes. Positive pressure engages multi-channel immunostaining mode, while negative pressure switches to whole-chamber immunostaining mode. (iv) Working principle of Dumi and its comparison conventional immunostaining. Static incubation suffers from a surface depletion zone that limits binding. Dumi’s advection-driven flow maintains antibody concentration at the tissue interface, improving uniform delivery and binding kinetics. (**C**) Multi-channel immunostaining: (v) Simultaneous detection of 16 biomarkers is obtained in a single staining cycle. (vi) Staining intensity across all microchannels shows consistent expression levels. (**D**) Whole-chamber immunostaining: (vii) TSA-based mIF is achieved through multiple staining cycles (cycle count dictated by dye number, imaging capability, and tissue quality), enabling one-shot multiplexed imaging. (viii) Single-cell level mIF data supports diversity and correlation analysis of cellular phenotypes within the TME.

The first mode is multi-channel immunostaining. When external positive pressure is applied, the increased internal pressure within the air valve causes the detection layer to tightly seal against the tissue section surface, preventing leakage between adjacent channels. The 16 microchannels enable parallel staining of different biomarkers. Specific primary antibody solutions can be added to the open-loading reagent reservoirs, while additional reagents such as PBS buffer, blocking solution, and secondary antibody working solution are injected via the reagent feed inlet port of the 3D-printed manifold. The reagents from the upper 3D-printed manifold are then injected through the ports of the PDMS chip into the 16 microchannels, resulting in the immunostaining effect on the tissue section as shown in Fig. 1C(v). Quantitative analysis of the staining intensity within each microchannel allows for precise biomarker information and expression levels (Fig. 1C(vi)). We verified the crosstalk and uniformity (± 5%) characterization of the multi-channel IHC staining using tonsil samples (Fig. S4).

The second mode is whole-chamber immunostaining. When external negative pressure is applied, the volume compression inside the air valve causes the PDMS membrane at the top of the staining layer to bulge upwards. This separates the microchannel walls from the tissue, forming a large single chamber over the tissue section surface. In this mode, the entire tissue area can be stained simultaneously with a single biomarker. All reagents are loaded into open-loading reagent reservoirs, connected to a switching valve, and injected through the central reagent feed channels. TSA staining is performed multiple sequential cycles within Dumi until the full multiplex immunostaining process is complete. Based on the distinct emission spectra of the TSA dyes, we can effectively separate the signals corresponding to each biomarker (Fig. 1D(vii)) and perform correlation analysis of cells based on the mIF images (Fig. 1D(viii)).

### Optimization of Antibody Incubation Time, Flow Rate, and Antibody Concentration for Dumi

We evaluated antibody incubation at flow rates of 2, 4, and 6 μL min^−1^. Using a fixed primary antibody solution, we incubated sections at each flow rate for durations ranging from 5 to 30 min, and measured IHC staining intensity under each condition (Fig. S5). We selected human tonsil tissue as a model owing to its multiple, similarly structured germinal centers with relatively uniform cellular distribution, thereby approximating diffuse infiltration. Considering the variations in recommended dilution factors, antibody affinity, tissue penetration, and target abundance, we chose two primary antibodies representing high- and low-performance biomarkers and used their optimized staining parameters as benchmarks. We selected two primary antibodies with distinct characteristics: a CD20 monoclonal antibody targeting the membrane (representative images are shown in Fig. 2A, quantitative results in Fig. 2B); a Ki67 polyclonal antibody targeting the nuclear (representative images in Fig. 2C, quantitative results in Fig. 2D). Both primary antibodies were used at concentrations previously validated through conventional experiments (CD20 at 1:2000; Ki67 at 1:400). Two-way ANOVA analysis showed that both incubation time and flow rate significantly affected staining intensity, with a notable interaction indicating that the influence of flow rate varied depending on the incubation time (Table S1 and table S2).

**Fig. 2.**
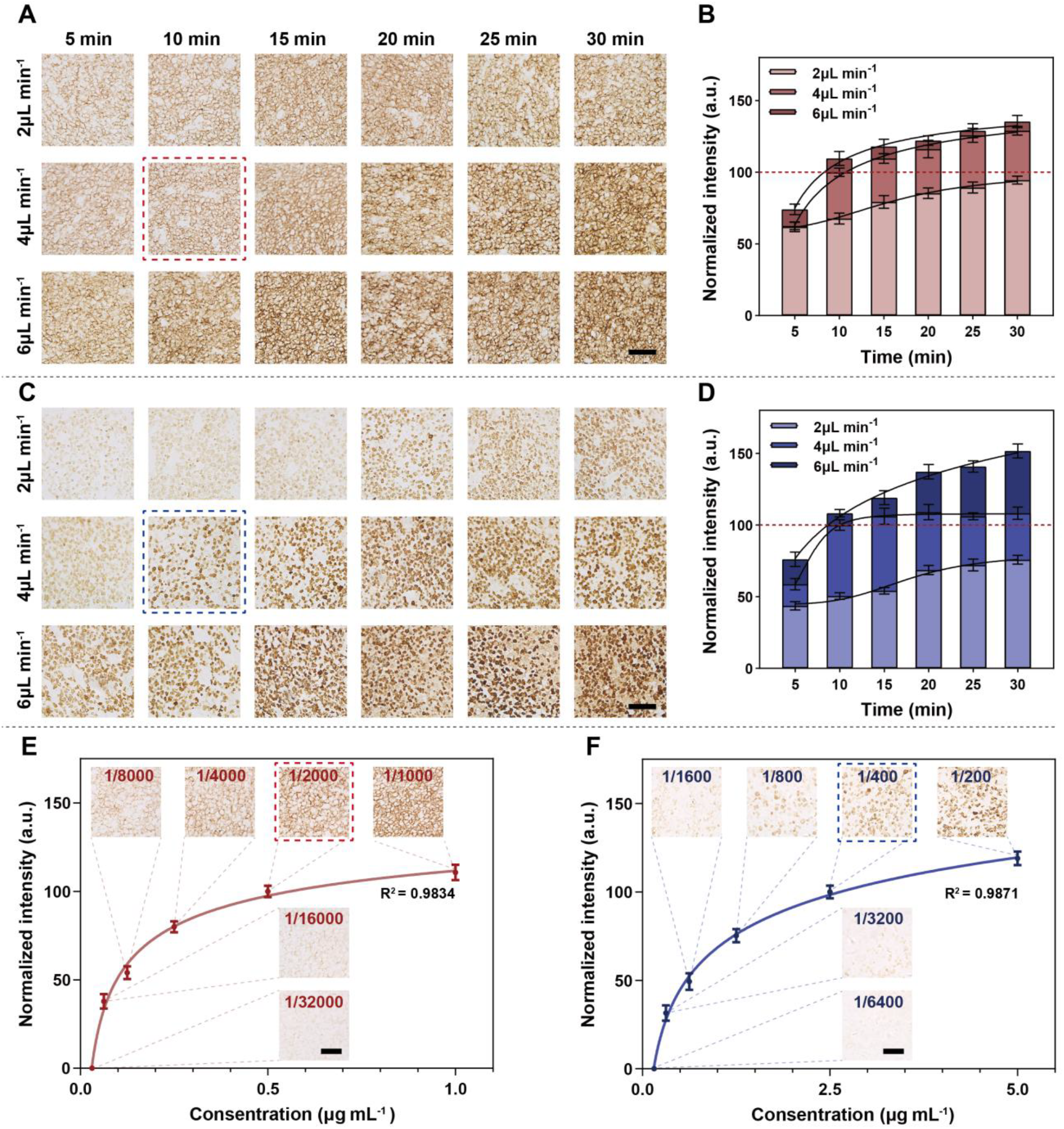
Optimization of flow rate, incubation time, and antibody concentration for Dumi-based immunostaining (Scale bars = 100 μm). (**A**) Representative images of CD20 staining in tonsil tissue at three flow rates over 5–30 min (n = 20). (**B**) Quantification of CD20 staining intensity over time at each flow rate. (**C**) Representative images of Ki67 staining in tonsil tissue at three flow rates over 5–30 min (n = 20). (**D**) Quantification of Ki67 staining intensity over time at each flow rate. (**E**) Fitted curve showing changes in staining intensity with varying CD20 concentrations. (**F**) Fitted curve showing changes in staining intensity with varying Ki67 concentrations.

Control over antigen-antibody reaction kinetics is crucial for quantitative IHC analysis. Incomplete antibody binding leads to underestimation of true antigen level, while oversaturation increases non-specific signals, both of which compromise quantitative accuracy. The 2 μL min^−1^ groups display a linear increase in staining intensity, indicating that saturation has not yet been reached. In contrast, staining intensities in the 4 μL min^−1^ and 6 μL min^−1^ groups plateau after 10 min, suggesting near-saturation, with any further gradual increases reflecting non-specific binding(Fig. 2, B and D). Examination of representative images and background signal quantification (Fig. S6) revealed a marked increase in non-specific background staining for Ki67 at the 6 μL min^−1^ group, indicating that higher flow rates exacerbate non-specific background. However, CD20 exhibited minimal background variation, potentially attributable to the high binding-site specificity of the monoclonal antibody. Based on these findings, at the antibody concentrations recommended for conventional IHC, we selected an incubation at 4 µL min^−1^ for 10 min to minimize non-specific background while conserving time and reagent (all subsequent antibody incubations were performed at 4 µL min^−1^). To further validate the suitability of this condition, we performed IHC staining for CD20 and Ki67 using a range of primary antibody concentrations (Fig. 2, E and F). The results showed that at concentrations of 1:2000 for CD20 and 1:400 for Ki67, antigen-antibody binding is approached saturation (reaching 92% and 84% of maximum observed intensity, respectively) without excessive non-specific signal. The similar shapes of the concentration-response curves for both antibodies further demonstrate the consistency and low variability of the Dumi-based method.

### Characterization and Validation of Dumi-based Whole-Chamber Immunostaining

We compared Dumi’s whole-chamber staining with conventional manual IHC by quantifying DAB intensity across equal-area ROIs. Tonsil sections stained for CD20 were used, with the primary antibody incubated at a flow rate of 4 μL min^−1^ for 10 min to evaluate staining uniformity across the chamber. These results were compared to those obtained from conventional manual staining.

Fig. 3A displays conventional IHC staining, performed manually by an experienced technician, with a primary antibody incubation time of 1.5 h. Fig. 3B shows Dumi-based whole-chamber IHC staining. A serial section adjacent to that in Fig. 3A was pre-treatment and placed directly into Dumi, completing the entire workflow from blocking to secondary antibody incubation in under 50 min (including a 10 min primary antibody incubation). To evaluate the uniformity differences, we segmented the stained regions from both the manual IHC and whole-chamber IHC images into 874 ROIs of consistent pixel size and measured the average DAB intensity for each ROI. Fig. 3C compares the staining intensity distributions across the 874 ROIs, showing highly similar distribution patterns between the two methods. Further Gaussian fitting (Fig. 3D) reveals that the data obtained from whole-chamber IHC show a better fit, indicating superior staining uniformity and overall performance. Bland-Altman analysis (Fig. 3E) demonstrates that the dispersion of the differences falls within an acceptable range (SD of bias = 0.03273), suggesting good agreement between the two methods in capturing the morphological distribution patterns of CD20 in the images. Quantitative analysis of IHC staining intensity data and the images themselves reveals that conventional IHC shows certain ROIs with higher intensity, visually corresponding to stronger staining at the tissue edges. This is likely an edge effect, possibly arising from higher local antibody concentrations at the periphery during static incubation.

**Fig. 3.**
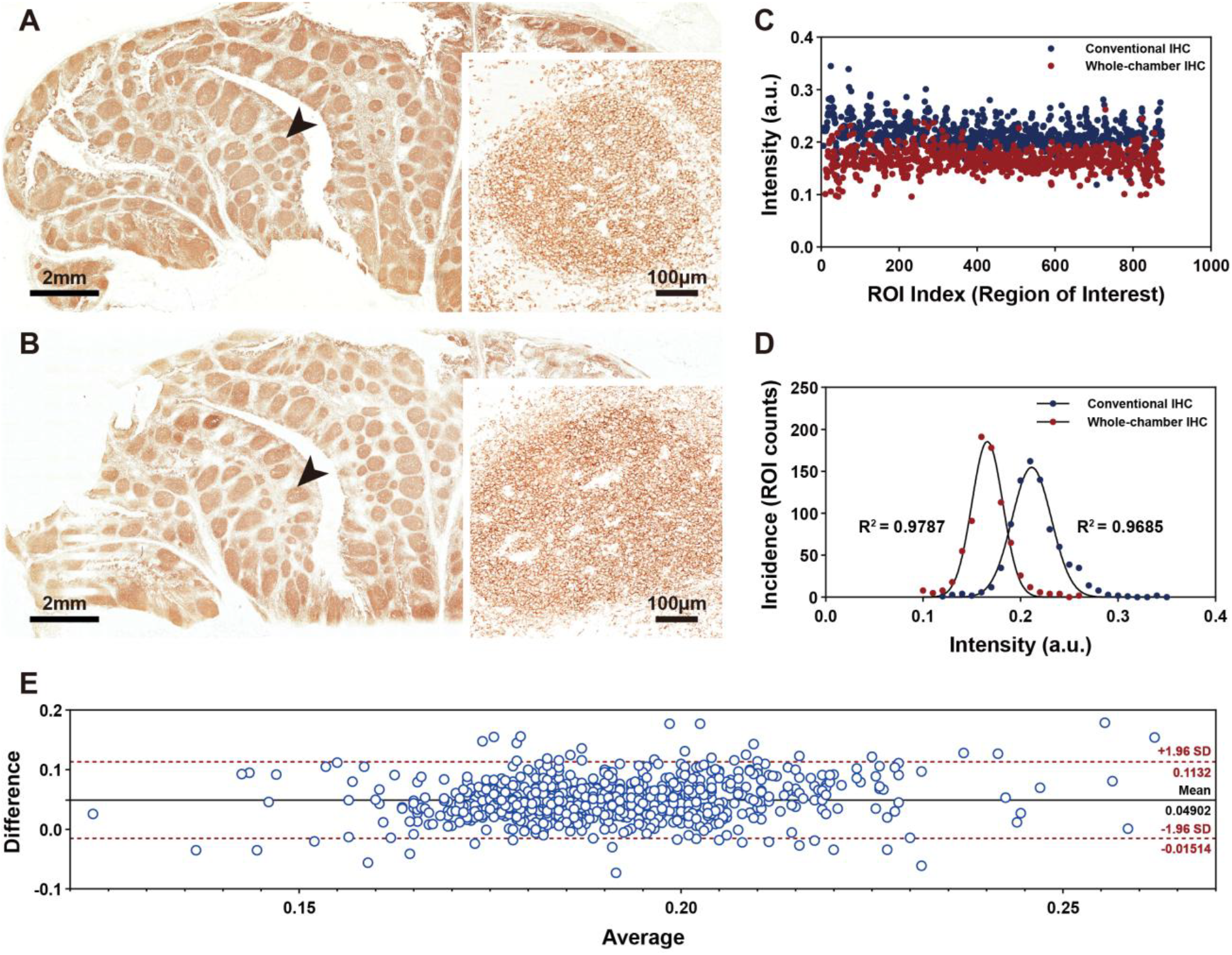
Comparative characterization of whole-chamber IHC and conventional IHC. (**A**) Conventional IHC result for CD20 staining in tonsil tissue. (**B**) Dumi-based whole-chamber IHC result for CD20 staining in tonsil tissue. (**C**) Both images were partitioned into 874 ROIs of consistent pixel size for intensity measurement. (**D**) Fitted Gaussian curve of the frequency distribution of staining intensity values from the both methods. (**E**) Bland-Altman analysis comparing the two staining methods. Red dashed lines indicate the upper and lower 95% limits of agreement.

### Dumi-based Biomarker Screening and TME Imaging Characterization

Following the validation of Dumi’s staining performance in whole-chamber mode, a six-cycle seven-color mIF experiment was performed using human tonsil sections. Tonsil tissue was chosen as a standard reference owing to its rich diversity of immune cell types and the distinct marker expression patterns between germinal centers and marginal zones, which facilitate assessment of staining accuracy (*25*). We selected different candidate biomarkers and performed mIHC staining separately within the 16 microchannels (Fig. S7, A and B). Based on the results of the mIHC staining, six biomarkers CD20, Ki-67, CD4, CD8a, CD68, and CD34 were selected for multiple cycles of Whole-chamber immunostaining. To achieve stable mIF images, we employed the TSA technique for mIF staining. However, due to the slow antibody incubation kinetics in conventional methods and the cycles required for TSA-based mIF staining, six-cycle staining typically takes 4-7 days. In contrast, each incubation step takes only 10 min within Dumi, enabling six-cycle mIF staining and imaging to be completed in one day. Fig. S7C shows the panoramic image of Dumi-based whole-chamber two-color IF staining on tonsil tissue. Fig. 4A shows a localized image of tonsil tissue stained with seven-plex IF, where immune cell lineages were marked by CD20 (B cells), CD4 (helper T cells), CD8 (cytotoxic T cells), and CD68 (macrophages), with Ki67 marking proliferating cells and CD34 marking endothelium cells. Image segmentation defined whole cells and their constituent nuclear and cytoplasm regions. We identified 7,047 cells, and quantified the expression of six biomarker fluorescence signals (Fig. 4B). We applied t-distributed Stochastic Neighbor Embedding (t-SNE) to map the seven-plex IF data (Fig. 4C(i)), achieving effective marker-based separation (Fig. 4C(ii)) and successfully resolved the expected cell populations. Fig. 4D provides cell counts following classification based on fluorescence signal intensity thresholds, with ‘Unknown’ referring to cells negative for all six biomarkers after excluding abnormal marker combinations. The most abundant cell population identified was proliferating B cells (Ki67^+^/CD20^+^), followed by cytotoxic T cells (CD8a^+^). Throughout the six-cycle During the TSA-based mIF staining process, Dumi demonstrated strong repeatability, minimal channel crosstalk during signal acquisition, and consistently produced precise single-cell-level data.

**Fig. 4.**
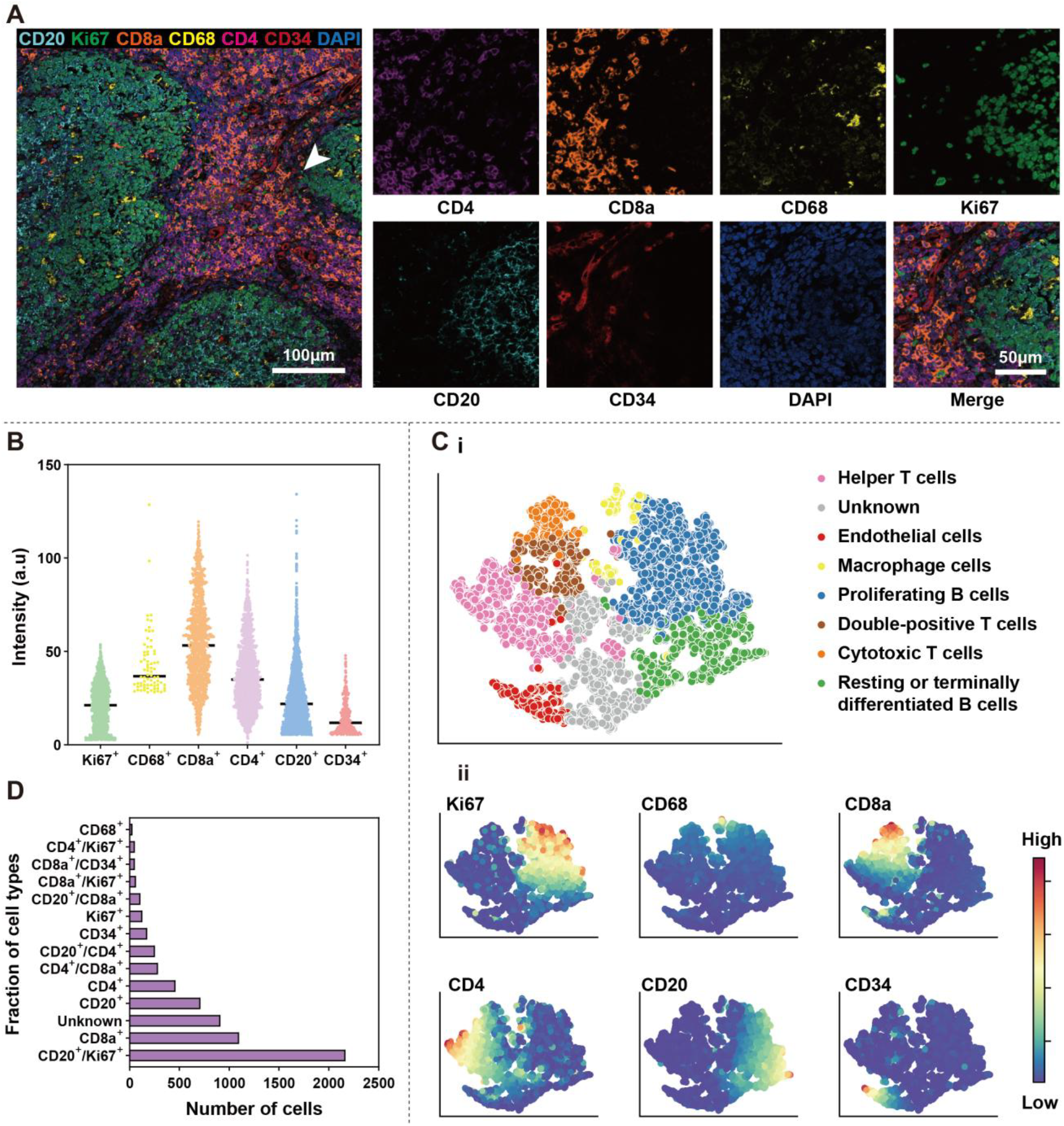
Execution and analysis of six-cycle, seven-color TSA-based mIF on tonsil tissue using Dumi. (**A**) Region of Interest (ROI) and representative local images from seven-plex imaging of tonsil tissue. (**B**) Distribution of staining intensities for positive cells across all markers (horizontal lines denote means). (**C**) Cell clustering and biomarker expression after segmentation: (i) t-SNE plot of all segmented cells from the ROI, clustered by PhenoGraph and manually annotated into eight distinct populations. (ii)Heatmaps showing the expression intensities of six biomarkers overlaid onto the t-SNE plot, corresponding to the manually identified cell populations in (i). (**D**) Cell counts for each identified population.

### Rapid Diagnosis of Clinical Pathological Samples

To assess the clinical diagnostics potential of Dumi, we performed rapid multi-channel immunostaining on four DLBCL samples obtained from different patients and primary sites (three sections for each case), including three germinal center B-cell–like (GCB) cases and one case of PCNS-DLBCL. All sections exhibited diffuse tumor cell infiltration and were centrally aligned beneath the microfluidic chip. Approximately 70% of the channels exhibited uniform marker density and staining intensity, whereas some regions showed variability in cell count and staining intensity across different fields of view. Consequently, we randomly selected fields of view within each channel and quantified intra-channel heterogeneity by calculating the standard deviation of staining intensity. Multi-channel mIHC staining results for diagnostic biomarkers in the PCNS-DLBCL case are shown in Fig. 5, A–C, with PBS (no primary antibody) microchannels served as negative controls. All biomarkers yielded clear DAB signals and enabled quantitative assessment of expression levels. Additional results from the other three cases are provided in Fig. S8–S10. The mIHC staining results of the four DLBCL cases were diagnosed by pathologists, and the biomarker expression levels for each case are shown in Fig. 5D, Fig. S8D, Fig. S9D, and Fig. S10D. We employed an immunophenotypic panel equivalent to that used in clinical reports. CD20, CD79A, and CD45 were used to confirm the B-cell origin of tumor cells, while CD3 and CD5 excluded T-cell lineage or special subtypes. CD10 and Bcl-6 served as germinal center markers, while MUM1 indicated activated B-cell status and was used to distinguish GCB from activated B-cell–like (ABC) subtypes (*26-28*). Bcl-2 and c-Myc were evaluated as indicators of tumor aggressiveness and prognosis, and Ki67 assessed proliferative activity (*29-31*). Given that PCNS-DLBCL often mimics other CNS tumors (e.g., gliomas, metastatic tumors) (*32, 33*), we included CD34 to assess vascular proliferation and heterogeneity (*34, 35*), and vimentin to rule to epithelial origin and indicate mesenchymal traits. Our single-slide Dumi -based diagnosis, the PCNS-DLBCL sample was classified as ABC subtype, DLBCL1 and DLBCL2 as GCB subtype (Fig. S8D and Fig. S9D), and DLBCL3 as GCB subtype with aberrant CD5 expression (Fig. S10D).(*36, 37*) Fig. 5E shows a heatmap of standardized biomarker expression differences across the four cases. All cases showed high levels of CD45 and Ki67, while expression levels of B-cell biomarkers CD20 and CD79A varied, reflecting underlying anatomical and pathological heterogeneity.

**Fig. 5.**
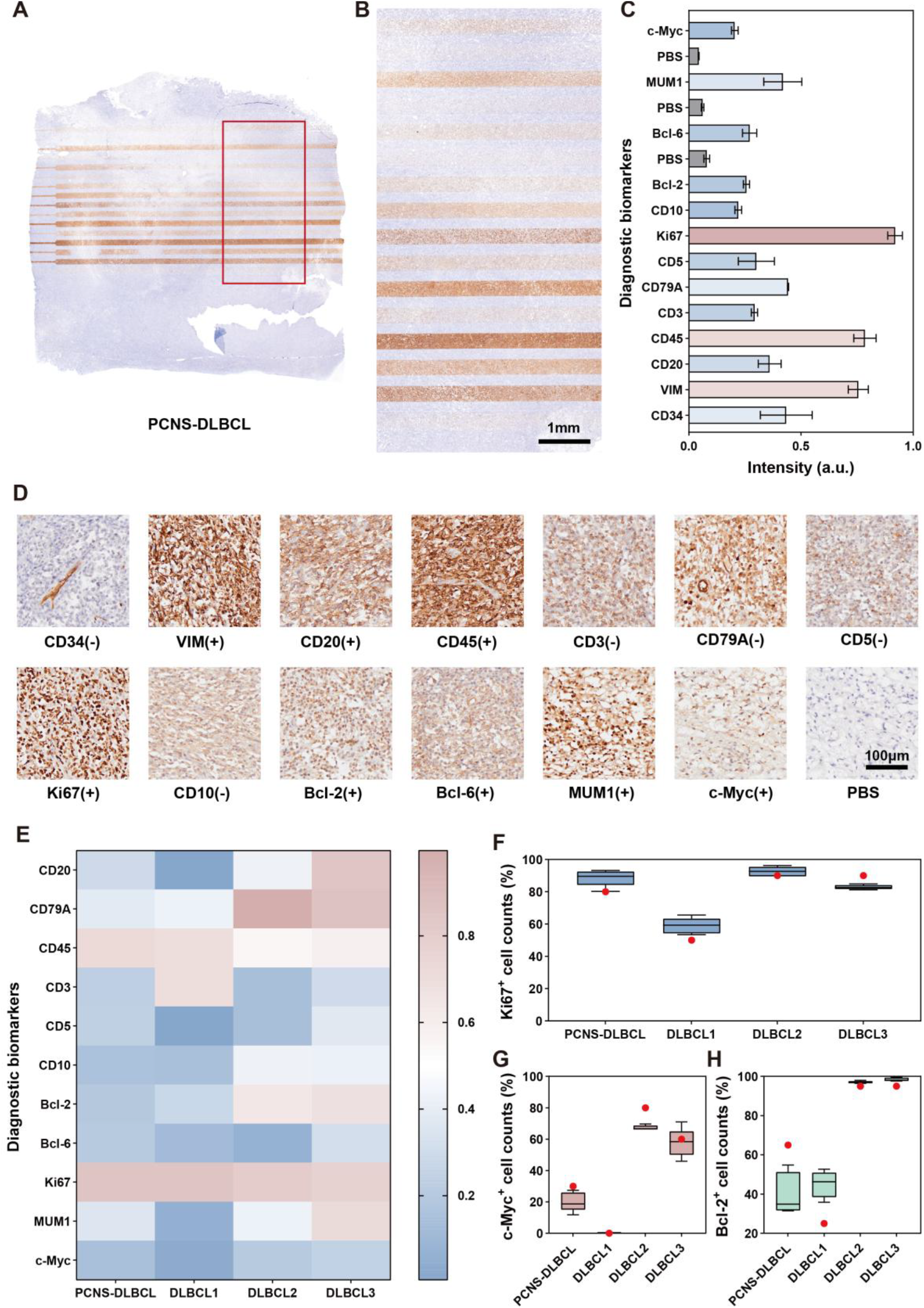
Rapid diagnosis analysis of four clinical DLBCL case using Dumi, including one PCNS-DLBCL case, and three others derived from spleen derived (DLBCL1), maxillary sinus derived (DLBCL2), and small intestine derived (DLBCL3). (**A**) Multi-channel IHC image of the PCNS-DLBCL case obtained using Dumi. (**B**) Magnified view of the region highlighted by the red box in (A). (**C**) Microchannel assignments and corresponding biomarkers overall expression levels corresponding sequentially to the 16 channels from top to bottom in image (B). The standard deviation represents different positions within each channel (n = 5). (**D**) Representative images of each biomarker in the PCNS-DLBCL case, along with pathologist-based diagnostic interpretations. ‘+’ denotes positive expression in tumor cells; ‘−’ denotes negative expression in tumor cells. (**E**) Heatmap showing the overall expression levels of each biomarker across all four cases. The scale column represents the normalized staining intensity. (**F-H**) Positive cell counts for Ki67, c-Myc, and Bcl-6 across the four clinical samples. Red dots represent the percentage estimates determined by pathologists. The standard deviation represents different positions within each channel (n = 5).

We next quantified key biomarkers relevant to clinical risk stratification and personalized treatment decisions by counting positive cells within high-cell-density “hotspot” regions of each channel, in accordance with standard IHC diagnostic practice. As shown in Fig. 5F, the counts of Ki67^+^ cells were assessed in four cases, where a high proliferation index (≥80%) was strongly correlated with poor overall survival (OS) (*38*). Fig. 5G presents the counts of c-Myc^+^ cells to assess oncogenic proliferative activity in each case, where ≥50% expression in IHC analysis may indicate gene rearrangement. Fig. 5H shows the counts of Bcl-2^+^ cells, reflecting anti-apoptotic capacity, where ≥50% of positivity is considered clinically significant. Co-expression of c-Myc and Bcl-2 in DLBCL is often associated with poor prognosis under conventional immunochemotherapy (*39*). Red dots represent percentage estimates annotated by pathologists based on visual inspection (empirical estimation). Quantitative results for Ki67^+^ and c-Myc^+^ cells (Fig. 5, F and G) showed good agreement with clinical diagnoses, while the slight discrepancy in Bcl-2 quantification (Fig. 5H) may be attributed to limitations in software-based cell type identification. These findings demonstrate the robust diagnostic and analytical capabilities of Dumi under resource-constrained conditions. However, given the limited number of available tissue sections, broader implementation of Dumi will require validation across additional patient cohorts.

### Construction of the TME in Clinical Pathological Samples

Building upon rapid subtype diagnosis of the PCNS-DLBCL case, this sample is ABC subtype, driven by persistent activation of the BCR/NF-κB pathway and multiple immune-evasion mechanisms (*40, 41*). To display the spatial distribution of various cell populations within PCNS-DLBCL relates to patient prognosis, we utilized the remaining slide after subtyping diagnosis to perform sequential whole-chamber TSA-based mIF staining, thereby enabling a deeper exploration of highly immunosuppressive TME of PCNS-DLBCL. Guided by tumor cell phenotypes and biomarker expression patterns, we selected CD20 to delineate tumor B cells, GFAP for astrocytes, CD34 for endothelial cells, and CD4, CD8a, and CD68 for immune cell lineage, thereby constructing a spatial TME map. Fig. 6A show the stitched image of the stained local tissue section, the selected ROI, and representative cells images within the ROI. Single-cell fluorescence intensity data extracted from the ROI were clustered using the PhenoGraph method, enabling the identification of cell populations including cytotoxic T cells, helper T cells, macrophages, and tumor B cells. The clustering results were visualized using t-SNE mapping (Fig. 6B(i)) (*42*). Hierarchical clustering illustrated similarities in biomarker expression across the identified clusters (Fig. 6B(ii)). Manually labels are shown on the left, and the histogram on the right shows the count of cells within each cluster, providing an intuitive understanding of the identified cell populations.

**Fig. 6.**
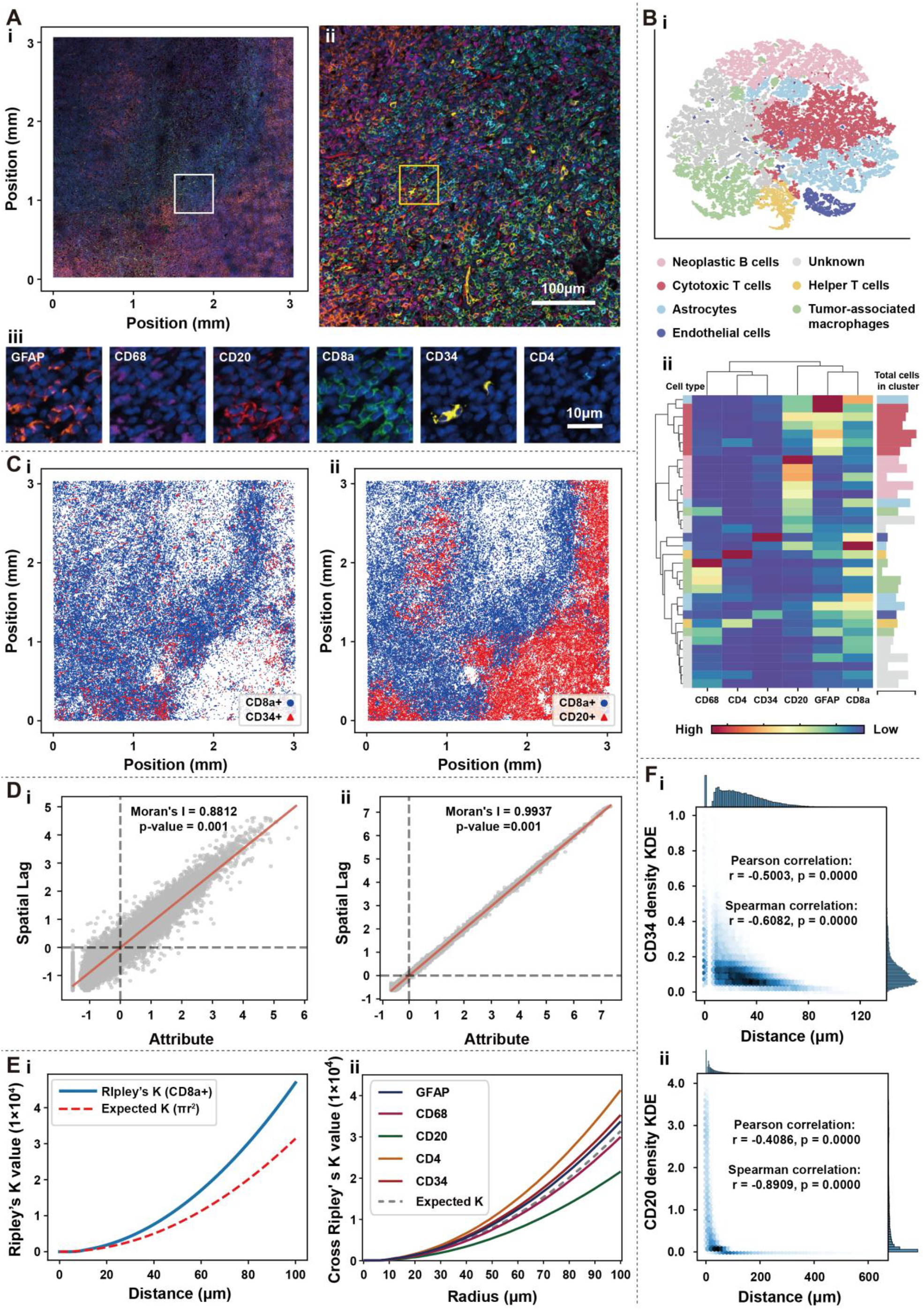
TSA-based mIF imaging and analysis of a PCNS-DLBCL sample processed by Dumi. (**A**) Stitched image of one tissue region containing 85,594 segmented cells (3,028.125 μm × 3,042 μm): (i) Overview with white box indicating the first zoom area. (ii) Zoomed-in view of the white-boxed region, with yellow box marking the second zoom. (iii) Single-channel view of the yellow-boxed area. (**B**) PhenoGraph-based cell clustering: (i) t-SNE plot of all segmented cells, annotated with manually inferred cell subpopulations. (ii) Hierarchical clustering dendrogram illustrating inter-cluster similarity based on marker expression. Heatmap colors represent biomarker expression levels. Annotation colors on the left shows cell populations, while the histogram on the right displays the number of cells within each cluster. (**C**) Spatial distribution maps: (i) CD8a^+^ cells relative to CD34^+^ cells. (ii) CD8a^+^ cells relative to CD20^+^ cells. (**D**) Moran’s I spatial autocorrelation analysis nearest neighbor distances: (i) CD8a^+^ to CD34^+^ cells. (ii) CD8a^+^ to CD20^+^ cells. (**E**) Ripley’s K analysis (within 100 μm): (i) Univariate Ripley’s K curve for CD8a^+^ cells. (ii) Cross Ripley’s K curves between CD8a^+^ cells and other cell populations (p = 0.005). (**F**) Joint distribution of nearest neighbor distance and KDE-estimated local cell density: (i) The top histogram: frequency distribution of CD8a^+^ to CD34^+^ cells. Right histogram: local CD34^+^ cell density. (ii) Joint distribution plot of CD8a^+^ to CD20^+^ cells distances and local CD20^+^ cell density.

Using centroid coordinates, we mapped CD8a^+^ cells within the tissue section image (Fig. 6A) to investigate their spatial relationships with other cell populations. We reconstructed spatial distribution maps of CD8a^+^ cells relative to each population (Fig. 6C(i) and 6C(ii) and Fig. S11). Next, we calculated the nearest neighbor distances from CD8a^+^ cells to each cell population (Fig. S12A–E) and applied Moran’s I index for spatial autocorrelation to assess overall clustering trends. The nearest neighbor distances from CD8a^+^ cells to CD34^+^ cells mostly aligned with the global regression line (red), indicating significant spatial aggregation of similarly spaced cells (Fig. 6D(i)). Distances to CD20^+^ cells exhibited high positive autocorrelation, reflecting an orderly, homogeneous distribution of CD8a^+^ cells around densely dispersed tumor cells (Fig. 6D(ii)). Distances to CD4^+^ cells showed a regular alignment (Fig. S12F), whereas those to CD68^+^ cells appeared more dispersed, suggesting a less uniform spatial relationship potentially influenced by local microenvironmental factors (Fig. S12G). Distances to GFAP^+^ cells showed only moderate positive autocorrelation, indicating a more random spatial relationship with glial or stromal elements (Fig. S12H). To further examine multiscale clustering behavior, we performed Ripley’s K analysis within 100 μm (*43*). Ripley’s K analysis of CD8a^+^ cells rose above the expectation under complete spatial randomness beyond 10 μm, indicating persisted clustering as spatial scale increased (Fig. 6E(i)). We then computed cross-Ripley’s K functions between CD8a^+^ cells and CD20^+^, CD34^+^, CD4^+^, CD68^+^, and GFAP^+^ cells (Fig. 6E(ii)). Results showed that within 100 μm, CD4^+^, CD34^+^, and GFAP^+^ cells tend to cluster with CD8a^+^ cells, whereas CD20^+^ cells exhibit clear spatial exclusion, and CD68^+^ cells showed a slight exclusion pattern.

To further visualize the association between CD8a^+^ cell enrichment and other cell populations, we used Kernel Density Estimation (KDE) to calculate the local densities of target cells at the locations of CD8a^+^ cell (Fig. S12I–K). As shown in Fig. 6F(i), CD8a^+^ cell positioning appeared to be influenced by local vascular density, with 87.4% of CD8a^+^ cells located within 50 μm of vessels (Fig. S12A), consistent with previous reports suggesting that high perivascular infiltration facilitates migration into the tumor core (*44*). Fig. 6F(ii) showed that many CD8a^+^ cells were adjacent to tumor cells, implying potential immune recognition and cytotoxic T cell recruitment. However, such proximity might also reflect local immune suppression or escape mechanisms (*45*). Therefore, integrating functional markers of CD8a^+^ cells (e.g., activation or exhaustion biomarkers) would be necessary to refine their prognostic significance. Additionally, CD8a^+^ cells selectively near and concentrate in regions enriched for CD4^+^ cells, CD68^+^ cells, and GFAP^+^ cells. Notably, the most pronounced monotonic co-enrichment occurs with CD4^+^ cells (Fig. S12I), the nearest-neighbor distance peak of 2–15 μm to CD68^+^ cells suggests a zone of efficient immune surveillance and clearance (Fig. S12J), and GFAP^+^ cells exhibit the strongest linear co-enrichment (Fig. S12K). These tight spatial associations suggest that CD8a^+^ cells may have a synergistic effect or mutual regulation with these target populations. The standardized outputs generated by Dumi, including cell coordinates, density metrics, and subpopulation classifications, provided a reproducible foundation for TME analyses, while deeper mechanistic interpretation will require expert-driven exploration.

## Discussion

In this study, we addressing the challenge of reconciling complex pathological subtyping demands with limited tissue availability, particularly in diffusely infiltrative rare tumors such as PCNS-DLBCL, by developing Dumi, a fully integrated platform for histopathological sections. Dumi enables seamless progression from rapid diagnostic biomarkers screening to high-precision spatial analysis of the TME. Utilizing a sequentially stacked 3D microfluidic chip design, we validated: 1) two operational modes of Dumi and the robustness of each detection microchannel; 2) optimization of incubation flow rates and durations, enabling precise control over staining time and conditions to improve the workflow reproducibility; 3) multiplexed detection of 16 diagnostic biomarkers on a single tissue section, reducing time and reagent consumption by over 80%; and 4) successful implementation of Dumi for six-cycle TSA-based mIF staining, enabling multidimensional, single-cell resolution analysis of TME. This platform empowers pathologists to rapidly obtain accurate in situ histological data. It serves as a bridge between diagnosis and prognostic evaluation, while alleviating the core issue of limited tissue availability in rare tumors studies. To validate Dumi’s clinical utility, we conducted diagnostic-level analyses on tissue samples from four DLBCL patients and obtained results comparable to conventional clinical IHC diagnoses. Furthermore, we performed a preliminary TME analysis on the seven-color stained PCNS-DLBCL sample. Experimental results confirmed that both operational modes of Dumi produce staining quality comparable to conventional immunostaining, while its precisely controlled fluid exchange process enables accurate adjustment of flow rates and incubation durations, resulting in significantly faster staining and enhanced reproducibility.

In summary, we have presented a microfluidic platform, Dumi, capable of performing both multi-channel and whole-chamber continuous immunostaining on a single slide. We detailed a complete workflow designed to extract comprehensive information from just 1-2 tissue sections, ranging from diagnostic subtyping to TME evaluation. Importantly, the potential applications of Dumi extend beyond this study. In the future, the platform could be adapted to support a broad range of tissue-based biochemical assays, including in situ hybridization, spatial transcriptomics, and genomics research. Integration of upstream processes such as deparaffinization, hydration, and heat-induced antigen retrieval could further streamline the workflow. Moreover, improvements in device architecture and materials may enable greater automation, thereby reducing labor demands. One limitation is that the DLBCL cases examined featured relatively homogeneous tumor cells distribution, which minimized the influence of tissue heterogeneity in multi-channel detection. Moreover, because diagnostic biopsy tissues are exceedingly scarce, we validated Dumi using surgically resected tumor samples; its tissue-saving advantage will be even more pronounced when applied to smaller biopsy tissues. For highly heterogeneous tissues, subsequent in-depth research will be carried out by incorporating prior morphological information, such as H&E staining, to guide ROI identification and optimize multi-channel assignment, while also guiding whole-chamber staining, thereby enhancing accuracy and broadening Dumi’s applicability. Overall, this work establishes a novel technical paradigm for rare tissue analysis, providing a methodological foundation for the clinical translation of integrated diagnostic-therapeutic systems and TME-driven personalized therapies.

## Materials and Methods

### Microfluidic chip fabrication and pretreatment for immunostaining

The 3D-printed fluidic exchange manifold and the microfluidic chip molds were fabricated using a projection micro-stereolithography (PμSL) based 3D printer (microArch S240, BMF Precision Technology, China) with HTL resin (Yellow-20, BMF Precision Technology, China) (Fig. S1). Post-printing treatments included active washing in 100% isopropanol, UV curing for 1h to eliminate residual resin and release internal stresses, followed by thermal treatment at 50°C for 2 h in an oven to prevent deformation. The 3D-printed fluidic exchange manifold was further soaked in 100% isopropanol overnight, then flushed with ultrapure water. For PDMS (RTV615A, Momentive, USA) replication, the 3D-printed microfluidic chip molds were repeatedly immersed in isopropanol to detoxify initiators prior to use (*46*). Air valve and biomarker staining layers were then molded and cured. Both layers were processed in a plasma cleaner (CPC-Eplus, CIF, China) with air at 10 Pa and 25 W for 60 s. After removal, the upper and lower layers were aligned with the ports and the air valve were positioned above the detection microchannels within 1 min, and then heated in an oven at 85°C for 20 min, to form a strong irreversible bond. Before the PDMS chips were used, they were processed in a plasma cleaner with 10 Pa, 25 W of air for 60 s. After removal, they were immersed in 75% isopropanol and cleaned with an ultrasonic machine (SN-QX-32D, Sunne, China) for 5 min, and then transferred to ultrasonic cleaning with ultrapure water for 5 min to enhance hydrophilicity and cleanliness of the microchannel surfaces.

### Microfluidic System for Rapid Immunostaining

The system was constructed using OEM components, including a positive pressure module (FU-FEZ-0100, Fluigent, France) and a negative pressure module (FU-FEZ-0800, Fluigent, France) controlled hydraulic pressure and pneumatic pressure. Switching valve (IVMSW1, Fluigent, France) regulated sequential reagent delivery, and two flow sensors (FLU-L+-OEM, Fluigent, France) monitored real-time fluid velocity changes at the chip interface. Computer-based fluid management, fully automated programming control, and real-time monitoring were integrated through a custom software interface. The 3D-printed fluidic exchange manifold, microfluidic chip, and tissue section achieved pressure-based reversible sealing via clamps and were immobilized on a customized microscope stage.

### FFPE Sample Preparation

The collection and use of human samples in this study were approved by the Biomedical Ethics Committee of Haikou People’s Hospital (Approval No. 2025-(Ethics)-073). Tonsils and lymphoma tissue samples preserved as paraffin blocks were provided by Haikou People’s Hospital and derived from archived pathological sections after completed clinical diagnostic reports had been issued. Prior to conventional staining and microfluidic-based immunostaining, all sections underwent dewaxing using eco-friendly dewaxing solution (YULU, China) and rehydration followed by a graded ethanol series. Antigen retrieval was performed with Tris-EDTA (pH = 9.0, G1218, Servicebio, China) in a microwave oven (MZC-2070MW, Haier, China) under the following protocol: medium power (8 min), standing (8 min), medium-low power (7 min), while preventing buffer evaporation and tissue drying. After natural cooling, samples were rinsed thoroughly with PBS buffer (RC20859, AMEKO, China).

### Conventional IHC Staining

Following antigen retrieval, control group sections were circumscribed with a histochemical pen and treated with 3% H_2_O_2_ for 25 min at room temperature in the dark to quench endogenous peroxidase activity. Sections were blocked with 3% BSA (GC305010, Servicebio, China) for 30 min to suppress nonspecific binding. After PBS rinsing, mouse-derived or rabbit-derived primary antibody dilution (table S3) were applied and incubated for 1 h in a wet box. HRP-conjugated secondary antibody dilution (SV0001/SV0002, Boster, China) matching the primary antibody species were then added for 30 min at room temperature. DAB chromogen (AR1027, Boster, China) was freshly prepared and applied. Sections were counterstained with hematoxylin (G1004, Servicebio, China) for 3 min, differentiated in hydrochloric acid-ethanol (R33067, Yuanye, China), and returned to blue by 1% ammonia (R20788, Yuanye, China). For dehydration and clearing, sections were sequentially immersed in 75% ethanol (5 min), 85% ethanol (5 min), anhydrous ethanol I (5 min), eco-friendly dewaxing solution I (5 min), eco-friendly dewaxing solution II (5 min). Air-dried sections were mounted with neutral balsam (S30509, Yuanye, China).

### Dumi-Based Rapid mIHC Staining

Following antigen retrieval, tissue sections were sequentially assembled on the microfluidic platform with the 3D-printed fluidic exchange manifold, microfluidic chip, and tissue section, enabling post-assay removal for standard dehydration, clearing, and mounting at room temperature. Additional reagents were delivered via a central reagent feed channels under positive pressure to parallel microchannels of PDMS chip and managed by a switching valve. Reagents that need to be specifically added are injected through the open-loading reagent reservoir, and fluid extraction was driven by a negative pressure pump connected to chip outlets. After assembly, negative pressure was applied to air valve to form a chamber for PBS rinsing between all steps, followed by sequential delivery of 3% H_2_O_2_ to block endogenous peroxidase and 3% BSA for nonspecific binding site blocking. Subsequently, positive pressure compressed biomarker staining layer onto the section for primary antibody incubation and HRP-secondary antibody binding, with post-PBS rinsing and freshly prepared DAB chromogen application under microscopic monitoring to control reaction duration. Upon completing all biomarkers, pressure pump were reset to zero, and the microfluidic chip disassembled to retrieve sections for dehydration, clearing, and mounting with neutral balsam.

### Dumi-Based Rapid TSA mIF Staining

Following antigen retrieval, tissue sections were loaded onto Dumi and subjected to the commercial TSA-mIF kit (G1257, Servicebio, China), which includes DAPI, IF440, IF488, IF546, IF594, IF647, IF700, Sudan Black solution, and anti-fade mounting medium. Negative pressure was applied to air valve to form a large chamber for PBS rinsing between all subsequent reagent steps. Sequential treatments included 3% H_2_O_2_ for endogenous peroxidase blocking and 3% BSA for nonspecific binding site blocking, both delivered at a flow rate of 4 μL min^−1^ optimized for antibody incubation efficacy. For multiplex immunostaining, sections were immersed in Tris-EDTA buffer (pH = 9.0) and microwave-reprocessed for antigen retrieval after each cycle. Six cycles were performed on Dumi, followed by DAPI nuclear counterstaining (5 min), Sudan Black treatment (5 min) to quench tissue autofluorescence, and mounting with anti-fade medium.

### IHC Image Acquisition and Analysis

All panoramic images were acquired using the slide scanner (SLIDEVIEW VS200, Olympus, Japan) under 10× and 20× objectives. For uniformity assessment, crosstalk elimination, and flow rate gradient analysis, IHC images used for staining intensity quantification were captured with a color camera (FL-20, TUCSEN, China) under a 40× objective with identical exposure time and LED light intensity. 10 random fields were selected within individual staining bands of each microchannel, and 5 fields with consistent cell density were analyzed using ImageJ software to quantify intensity. For hematoxylin counterstained nuclear samples, noise signals were separated via color deconvolution to isolate the DAB channel, followed by threshold filtering to remove background in non-stained areas, with mean optical density measured to quantify immunostaining intensity. For DAB images without hematoxylin counterstaining, images were converted to 8-bit format, processed with the same threshold filtering algorithm to eliminate background, and mean optical density was similarly calculated. In whole-chamber or conventional IHC staining, intensity values were quantified within a 19 × 46 matrix grid. For clinical samples, positive cell counting was performed in QuPath software, cells were first segmented via automated detection, manually classified as positive or negative through annotation, and the classifier was uniformly applied to all fields.

### IF Image Acquisition and Analysis

For TSA mIF samples, imaging was performed using the laser scanning confocal microscopy system (STELLARIS 5, Leica, Germany) with 10× and 20× objectives, equipped with a 405 nm laser and supercontinuum white light lasers (wavelength range: 485-685 nm), where each dye was excited at 100% peak intensity to minimize spectral crosstalk. Image analysis was conducted in QuPath software by first segmenting cells via automated detection, followed by classifier creation for each fluorescence channel using simple thresholding to categorize positive/negative cells. For clinical samples, classifiers were refined through machine learning-based training to enhance accuracy. Raw data including fluorescence intensity values and spatial coordinates of all cells were exported from QuPath for downstream statistical analysis in Python software. Intensity values of cellular markers were subjected to K-means clustering for tonsil samples and PhenoGraph clustering for clinical specimens, with manual annotation of inferred cell populations based on cluster features. Both clustering results were visualized using t-SNE dimensionality reduction. For clinical sample cluster heatmaps, hierarchical clustering was performed by calculating Euclidean distances between marker expression values across clusters, followed by Ward’s linkage method to iteratively merge clusters into a dendrogram. Spatial analysis utilized cell coordinate data, Ripley’s K analysis quantified cumulative CD8a+ cell counts within a 0–100 μm radius and compared them to the theoretical random distribution (πr^2^), while Moran’s I analysis employed a K-nearest neighbors (K = 5) weight matrix to assess spatial autocorrelation and interdependencies.

## Author contributions

Conceptualization: B.S., Z. H.

Methodology: Y. Z., Y. H., B.S., Z. H.

Investigation: Y. Z., Y. H., X. G., J. J.

Visualization: Y. Z., Y. H., X. G., J. J., F. X.

Supervision: B.S., Z. H.

Writing—original draft: Y. Z.,

Writing—review & editing: B.S., Z. H.

## Data and materials availability

All data needed to evaluate the conclusions in the paper are present in the paper and/or the Supplementary Materials.

## Supplementary Materials The PDF file includes

Supplementary Text

Figs. S1 to S12

Tables S1 to S4

