## supplementary materials for "Dual-Mode Microfluidic Immunostaining (Dumi) Device for Diagnostic Biomarkers Detection and Tumor Microenvironment Evaluation"

Yu Zhang *et al.*

**This PDF file includes:**

Supplementary Text
Figs. S1 to S12
Tables S1 to S4

**Supplementary Text**

Dumi workflow

Reversible Assembly of the Microfluidic Chip: Bonded PDMS chips and pretreated tonsil sections were sequentially placed into the recesses of the 3D-printed manifold, aligning all ports so that the microchannels contact the hydrated tissue surface. The assembly is fixed to the microscope stage using clamps and screws. Note that excessive negative pressure during fluid withdrawal can imprint the microchannels onto the tissue; to prevent this, we maintain suction below –100 mbar, use very low negative pressure during antibody incubation, and lift the biomarker staining layer via air valve during PBS washes to avoid PDMS contact and protect the tissue. This reversible assembly allows removal of the section for downstream processing and conventional slide mounting. All steps are performed at room temperature.

Reagent Loading: Sealed reservoirs connected to a switching valve were filled with 3% H2O2, 3% BSA, PBS buffer, and HRP-labeled goat anti-rabbit/mouse secondary antibody working solution.

Tissue Section and Microchannel Wetting: To maintain a fully hydrated environment and prevent air entrapment (which causes flow-rate disparities), PBS was injected into the 3D printed open-loading reagent reservoirs via positive pressure control module until all internal channels were filled. Without negative pressure, fluid preferentially filled the open-loading reservoirs connected to atmospheric pressure. Subsequent negative pressure at the chip outlet ensured complete PDMS microchannels filling and bubble elimination. A chamber formed over the tissue surface, immersing it in PBS.

Blocking Endogenous Peroxidase and Nonspecific Sites: After resetting negative pressure modules, 3% H2O2 was introduced via directional valve switching. A low negative pressure enabled slow reagent flow across the tissue for 10 min, followed by PBS rinsing and identical 3% BSA loading.

Section Unloading: After the secondary antibody incubation and PBS rinsing, tissue section were either disassembled for mounting or retained for workflows such as DAB chromogenic reaction or TSA dyes reaction and real-time observation of staining under the microscope.

Multiplexed Staining Cycles: If multiple rounds of immunostaining are required after one round is completed, such as TSA-based mIF experiments, antigen retrieval was performed to remove prior antibody complexes, followed by repeating the microfluidic workflow. The comparison of the time consumption between Dumi and conventional immunostaining is presented in table S1.

Crosstalk Characterization

In the 3D-printed fluidic exchange manifold we designed, the 16 open-loading reagent reservoirs can interconnect via the central reagent feed channels used for injecting additional reagents. To ensure that each open-loading reservoir can independently test different biomarkers, theoretically, we designed the inner wall of the circular channels with a 300 μm diameter to generate sufficient capillary force to counteract the hydrostatic pressure resulting from the open-loading reservoirs being connected to atmospheric pressure. This design also ensures that when liquid level differences exist between the open-loading reservoirs, the internal channels can withstand the mass transport caused by this pressure difference during the 1-5 min loading time, and furthermore, the time required for diffusion effects to reach detectable concentrations in adjacent channels is far greater than the time needed for the entire workflow. To further guarantee the absence of inter-channel crosstalk during the process of injecting from the open-loading reservoirs into the PDMS channels, we apply a small amount of positive pressure at the reagent feed inlet port. At this time, the central reagent feed channels still contains PBS buffer. The positive pressure pushes PBS towards the branch channels, forming a pressure barrier that separates the internal fluid paths of the individual reservoirs, preventing reagents in the reservoirs from diffusing or flowing back into the main channel. The flow sensor is used to monitor the flow rate in real-time for feedback regulation of the pressure control module, precisely maintaining the positive pressure at a preset low level. This avoids excessive positive pressure pushing too much PBS into the reservoirs, causing changes in the initial concentration, thus ensuring concentration accuracy.

To confirm the effectiveness of this method, we characterized the device for the absence of crosstalk between adjacent channels to ensure that the degree of diffusion between them does not affect the IHC results. On the human tonsil tissue, we performed staining for CD20 in alternating channels using the same concentration and experimental conditions. Specifically, during the primary antibody incubation step, CD20 antibody working solution was added to alternating open-loading reservoirs, while PBS was added to the reservoirs without antibody working solution as a control. The degree of staining crosstalk was verified in the channels where no antibody working solution was added. Fig. S4A shows the panoramic image of the IHC staining. Fig. S4B presents the results of directly measuring the staining intensity along the cross-section line in the Blue channel after converting the image to an RGB stack in ImageJ. Fig. S4C shows the quantitative results of staining intensity for each of the 16 channels separately. Visually, no obvious positive signal was observed in the channels loaded with PBS control. Furthermore, quantitative analysis could not automatically identify positive cells in the control channels using a thresholding algorithm due to their extremely low staining intensity. This result indicates that our device design theory and experimental procedure can provide an independent immunostaining environment for each microchannel, and the diffusion effect between reagents in the 3D-printed channels is negligible. This is an essential prerequisite for labeling and screening different antibodies in the 16 channels separately.

Uniformity Characterization

To demonstrate uniform fluidic velocity and independent biomarker detection environments across the 16 microchannels in our fabricated microfluidic chip, we performed multi-channel parallel IHC staining to compare inter-channel uniformity. Human tonsil tissue section with large area and structurally similar regions (containing multiple germinal centers) were selected to minimize tissue heterogeneity-induced variations in staining outcomes across channels. To validate staining uniformity, we applied identical antibody concentrations to all channels and compared IHC staining intensities. CD20 antibody (1:500 dilution) was selected due to the widespread distribution of B cells, enabling comprehensive characterization of staining consistency along the microchannels. All experimental parameters were strictly standardized, including antibody concentration, flow rate, and incubation time. Fluid was introduced into each channel via capillaries connected to a public reagent inlet, ensuring bubble-free flow within both 3D printed pipes and PDMS channels. The liquid level at the open-loading reservoirs was maintained above the 3D printed pipes to prevent flow obstruction or velocity disparities. Post-staining visualization with DAB enabled direct wet-state microscopic examination or post-processing and slide mounting. Fig. S3D shows panoramic images, Fig. S3E quantifies cross-sectional staining intensities in the blue channel after RGB stack conversion in ImageJ, and Fig. S3F displays quantified immunostaining intensities for all 16 channels. Results confirm that immunostaining conditions across the 16 microchannels are uniformly controlled by pneumatic pressure, with consistent staining intensity within individual channels. Well defined channel edges and absence of staining on the PDMS-contacted surfaces confirm no antibody leakage, ensuring fluid confinement to designated regions. By aligning tissue sections with specific chip positions, channels can be precisely targeted to anatomical structures or regions of interest, enabling biomarker localization adjustments based on prior histological knowledge in practical applications.

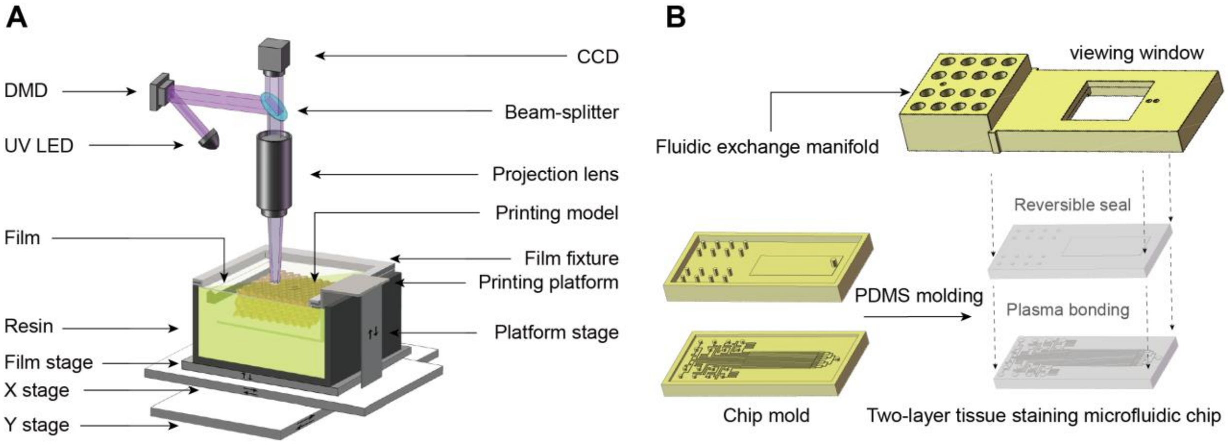


**Fig. S1. Fabrication of the 3D-printed fluidic exchange manifold and microfluidic chip molds based on projection micro stereolithography.** (**A**) Configuration and principle of the projection micro stereolithography system: ultraviolet light is projected onto the platform surface to additive manufacture the resin. (**B**) Comparison of two types of 3D-printed components and their functions: the fluidic exchange manifold features a recessed cavity at its bottom for accommodating the microfluidic chip and tissue section, with a viewing window directly above the tissue to enable real-time observation through the transparent PDMS under a microscope; the 3D-printed chip molds is used to pour and demold PDMS. After plasma bonding, the two layers form an integrated chip.

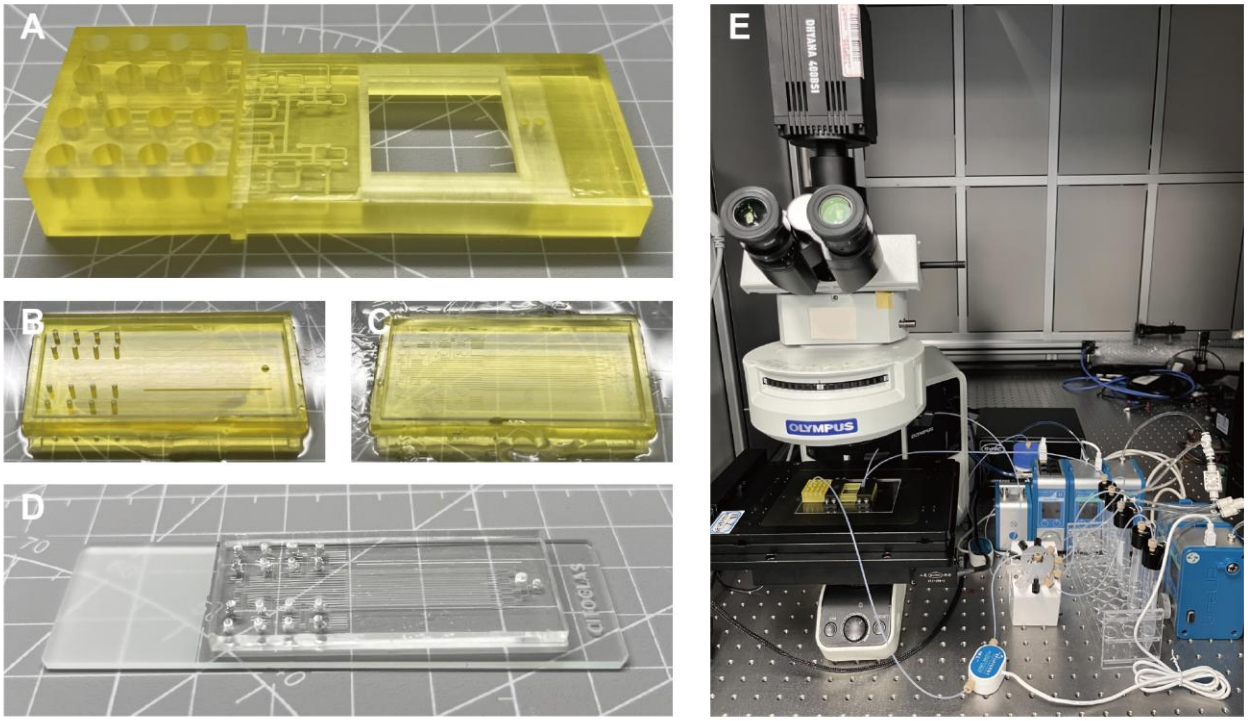


**Fig. S2. Photographs of the 3D-printed fluidic exchange manifold, the chip molds, and the complete Dumi microfluidic system.** (**A**) Photograph of the 3D-printed manifold for fluid exchange. (**B**) Photograph of the 3D-printed PDMS chip molds. (**C**) Photograph of the assembled Dumi integrated into the microscope stage.

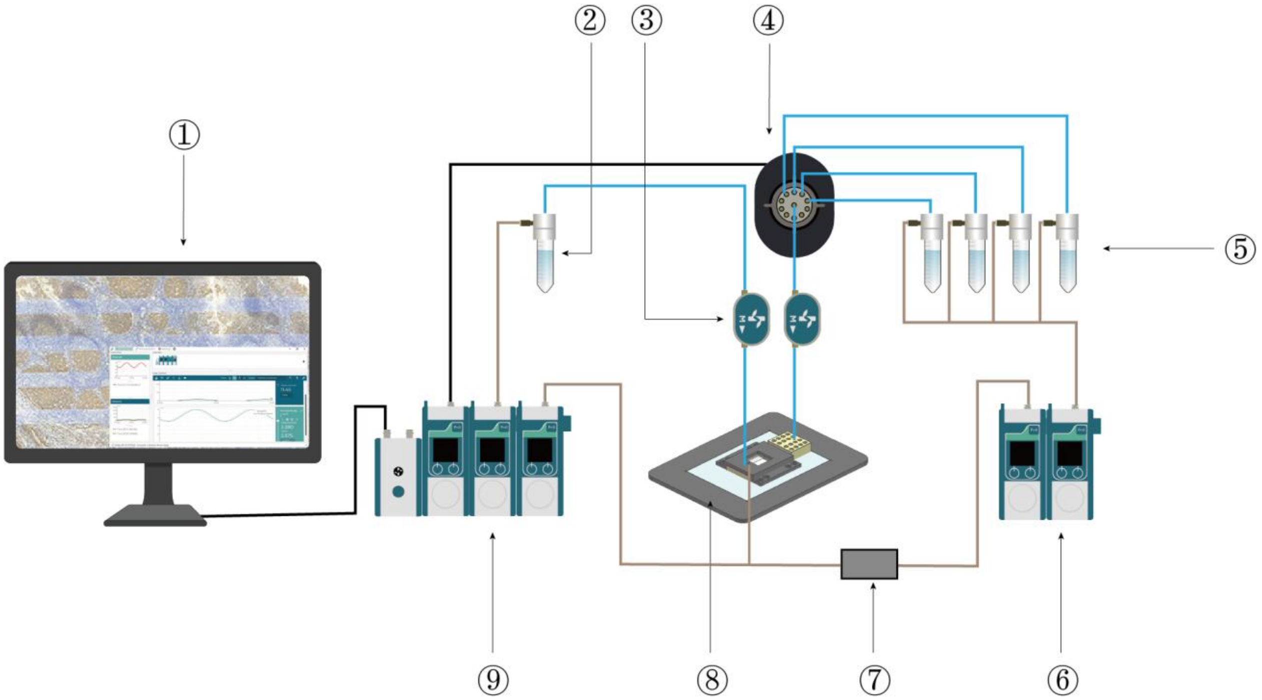


**Fig. S3. Schematic of the Dumi system's connection principles.** Includes: computer control unit (①), which sets pressure parameters, fluid-switching sequence, and timing via software; waste collection module (②) connected to the reagent outlet of the microfluidic chip (⑧) via a flow sensor (③); for fluid control, the positive pressure control module (⑥) supplies pressure to additonal reagents (⑤), the switching valve (④) selects the appropriate reagent and connects to the chip's reagent feet inlet port, and the negative pressure control module (⑨) draws fluid into the waste module (②); for air valve control, modules (⑥) and (⑨) connect to the chip's air intake via an air-switch (⑦) to inflate or deflate the valve.

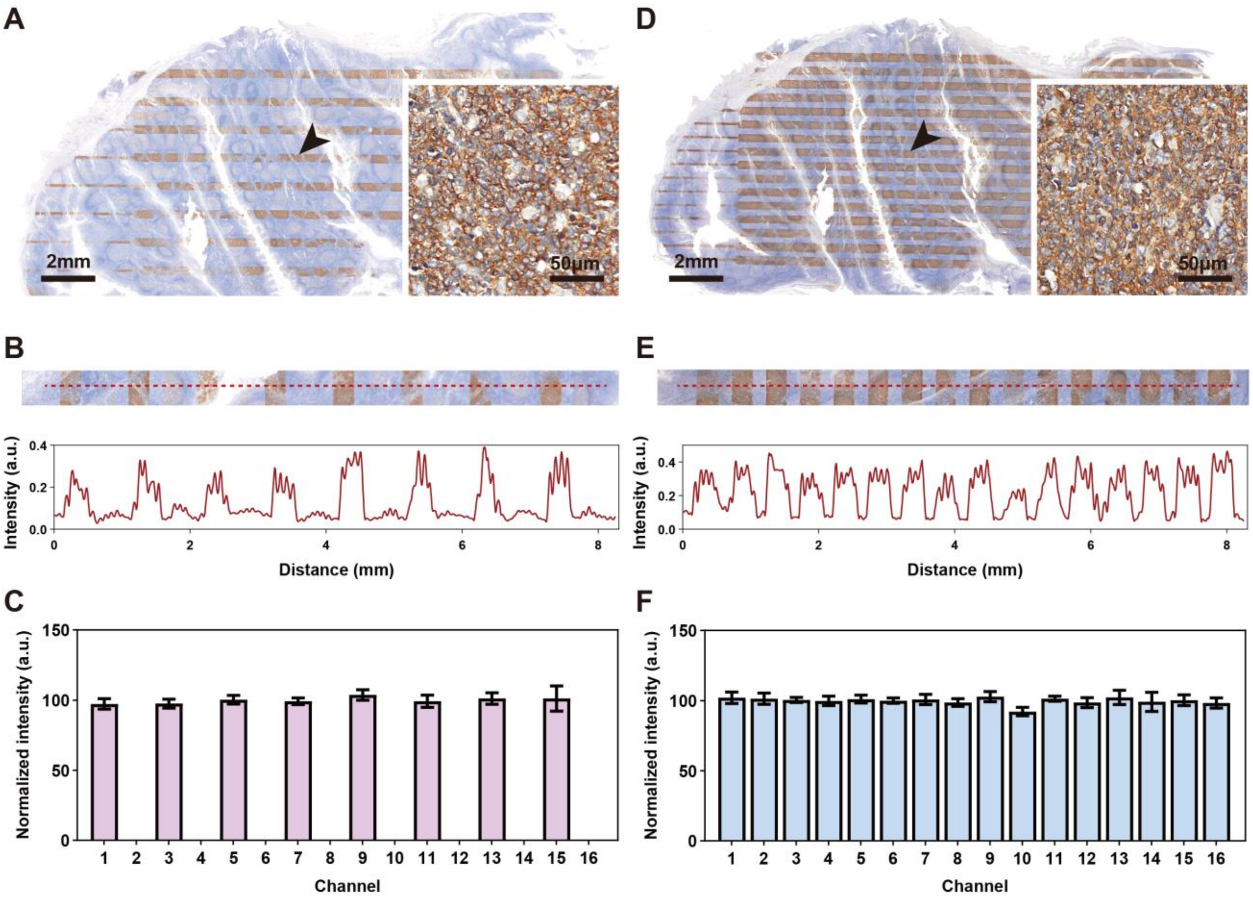


**Fig. S4. Characterization of crosstalk absence and uniformity.** (**A**) 8 alternating channels IHC staining was performed on tonsil tissue to verify whether crosstalk occurs between adjacent channels, and panoramic and detail images were obtained. Eight channels were stained for CD20 under identical experimental conditions. (**B**) Staining intensity profiles along the red dashed line drawn perpendicular to the channels in the panoramic image from (A). (**C**) Quantitative results of staining intensity for the 8 alternating channels during crosstalk absence characterization. Staining intensities were linearly normalized. The standard deviation represents different positions within each channel (n = 5). (**D**) multi-channel parallel IHC staining was performed to assess the uniformity of the 16 channels, and panoramic and detail images were obtained. All 16 channels were stained for CD20 under identical experimental conditions. (**E**) Staining intensity profiles along the red dashed line drawn perpendicular to the channels in the panoramic image from (D). (**F**) Quantitative results of staining intensity for the 16 channels during uniformity characterization. The standard deviation represents different positions within each channel (n = 5).

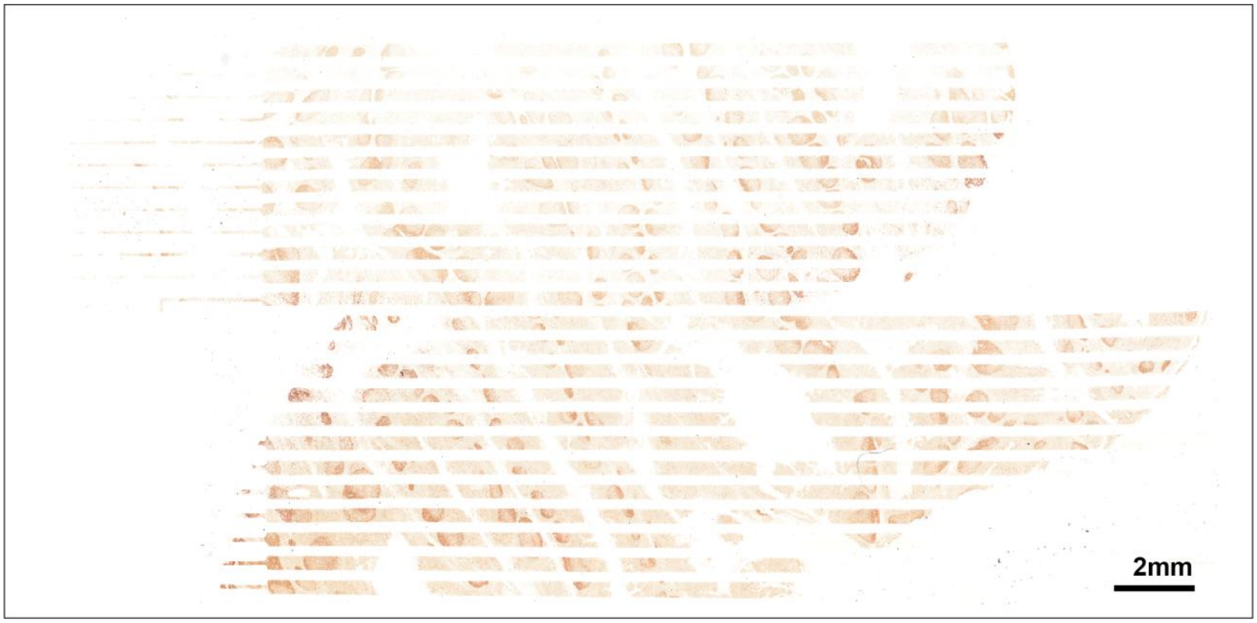


**Fig. S5. Panorama of parallel Ki67 gradient IHC staining across 16 channels at 4 µL min$^{-1}$ for 5–30 min.** Each group of 4 channels represents a parallel experiment at the same time point, and two slides were tiled.

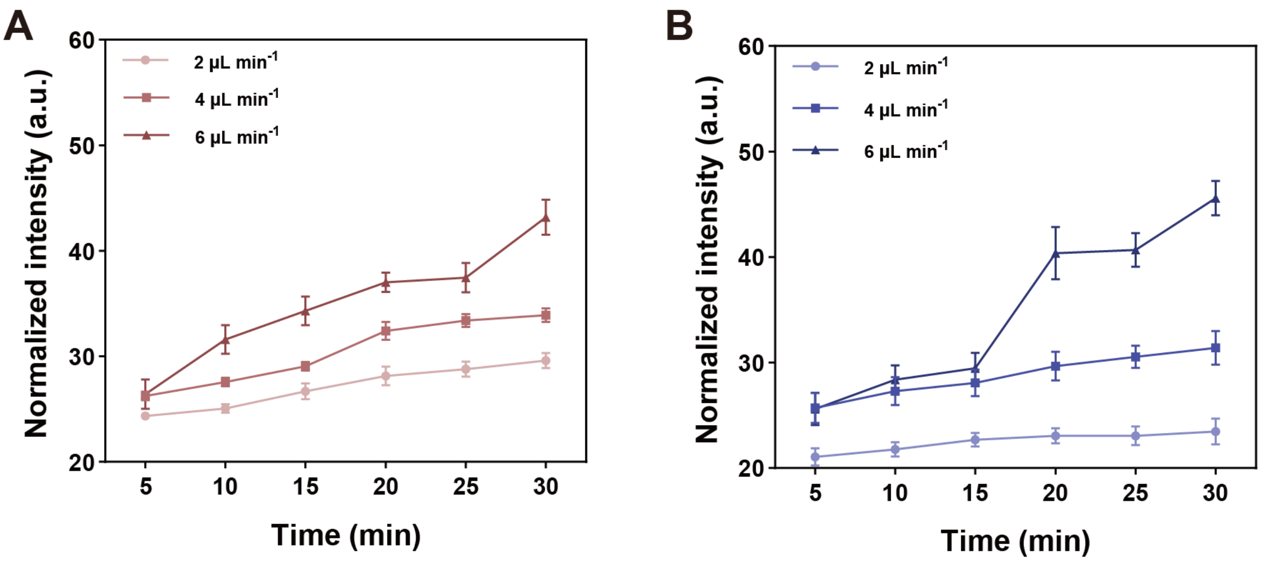


**Fig. S6. Nonspecific staining signal levels collected from the remaining areas after excluding regions positive for IHC staining, defined as background.** (**A**) Background signal levels for tonsil CD20 staining across different groups. (**B**) Background signal levels for tonsil Ki67 staining across different groups.

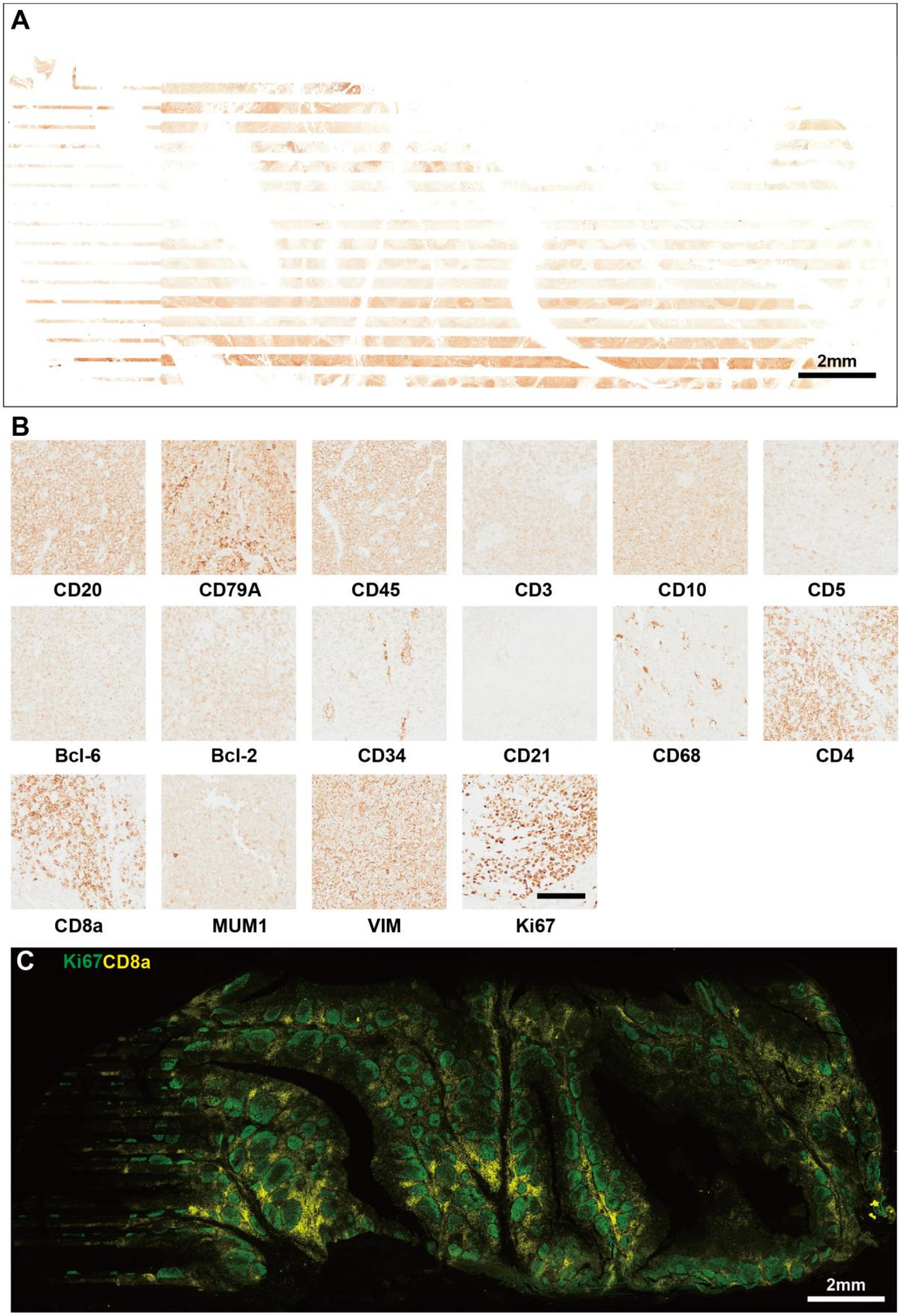
A
2mm
B
CD20
CD79A
CD45
CD3
CD10
CD5
Bcl-6
Bcl-2
CD34
CD21
CD68
CD4
CD8a
MUM1
VIM
Ki67
C
Ki67CD8a
2mm

**Fig. S7. Rapid screening of mIF biomarkers.** (**A**) Panorama of specific marker IHC staining in each of the 16 channels. (**B**) Names and detail images of the markers from top to bottom in (A) (scale bars = 100 μm). (**C**) Panorama after TSA-based mIF on tonsil tissue using the whole-chamber immunostaining mode, showing Ki67 and CD8a.

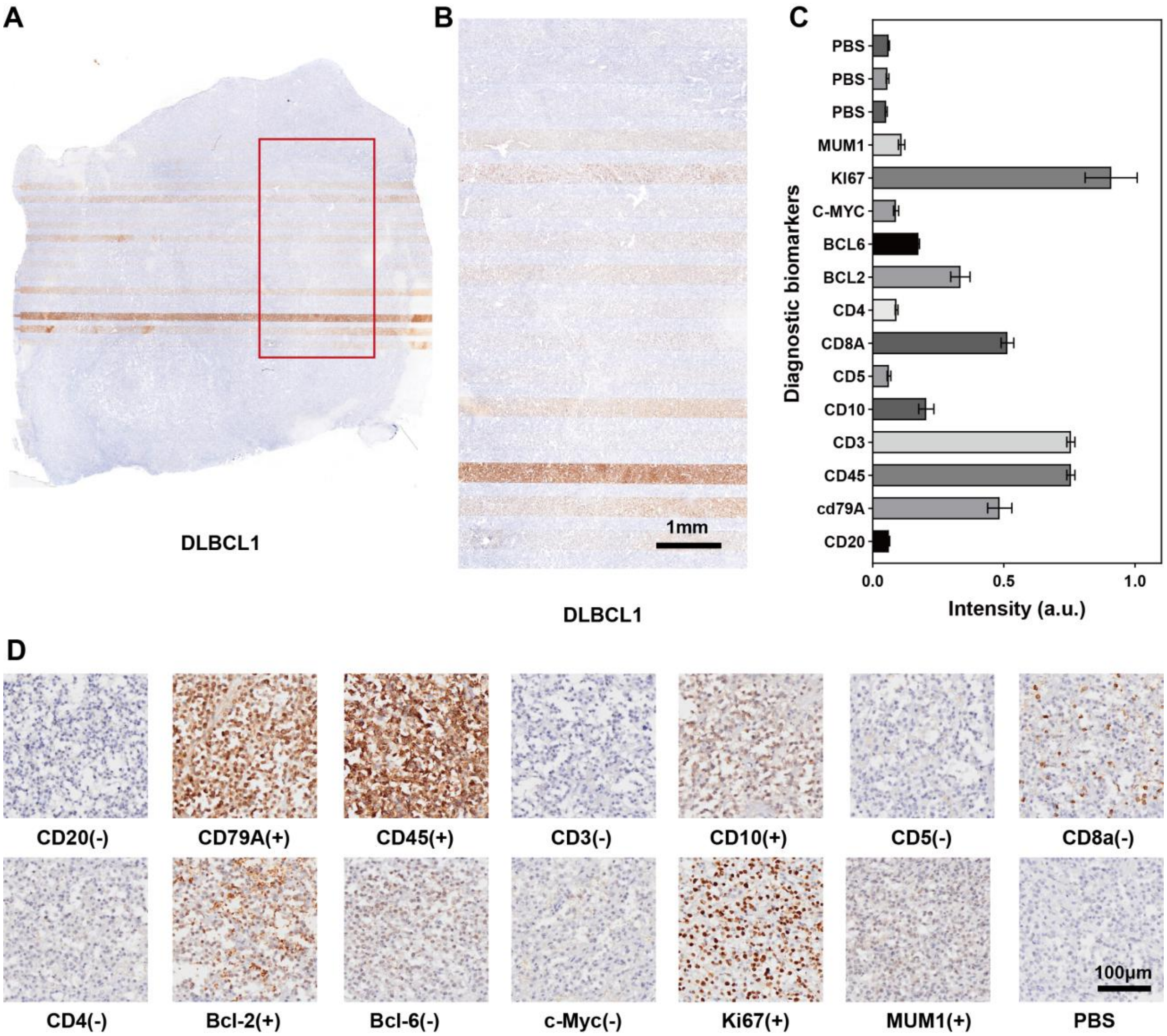


**Fig. S8. Single slide diagnostic markers expression for DLBCL1.** (**A**) DLBCL1 sample after Dumi-based multi-channel mIHC staining. (**B**) Magnified view of the mIHC image. **(C)** Markers corresponding to each channel in the image and their overall expression levels corresponding sequentially to the 16 channels from top to bottom in image (B). The standard deviation represents different positions within each channel (n = 5). (**D**) Representative images for each marker in the DLBCL1 sample and the corresponding diagnosis provided by pathologists based on these images. '+' denotes positivity in tumor cells; '−' denotes negativity in tumor cells.

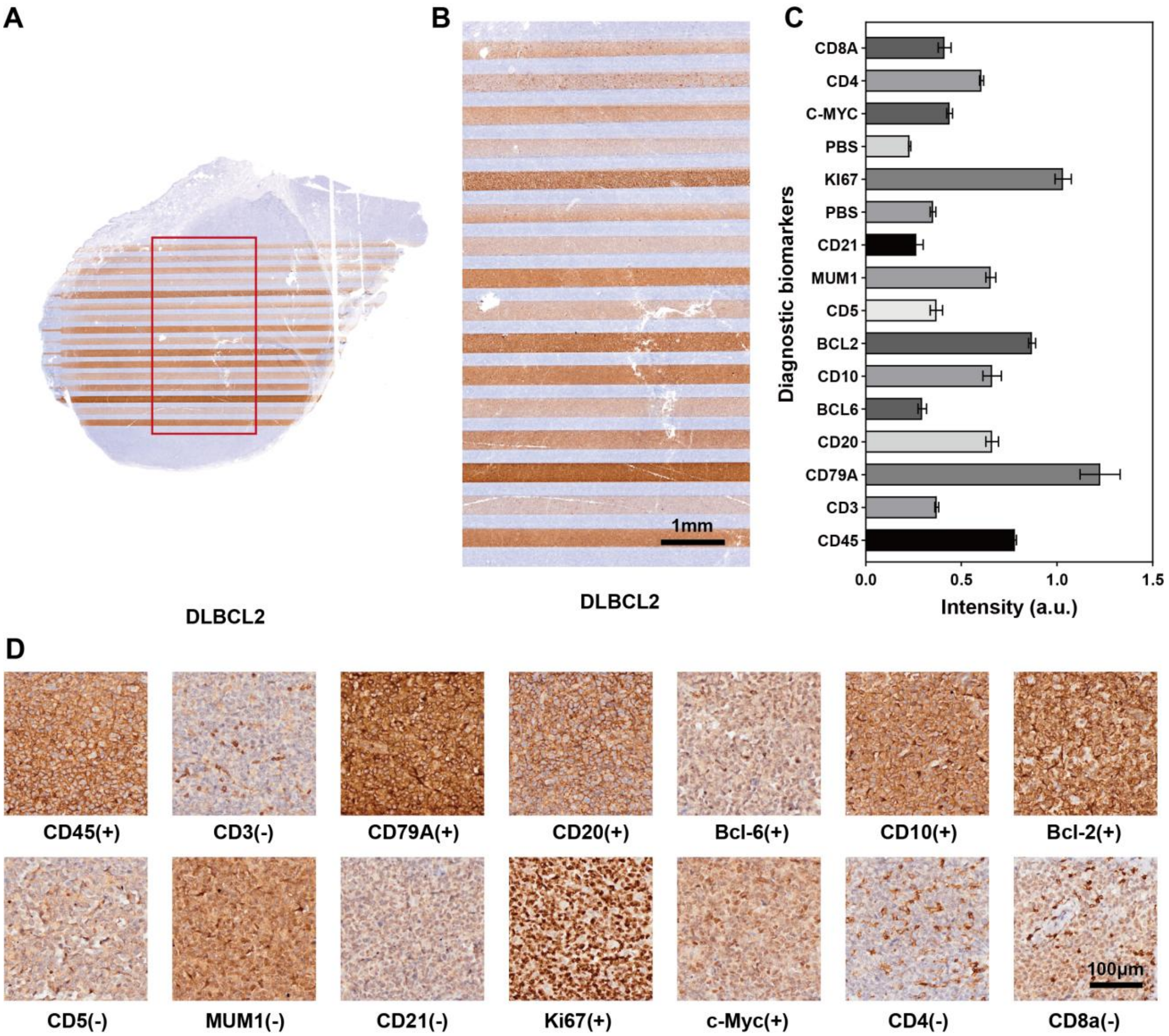


**Fig. S9. Single slide diagnostic markers expression for DLBCL2.** (**A**) DLBCL2 sample after Dumi-based multi-channel mIHC staining. (**B**) Magnified view of the mIHC image. (**C**) Markers corresponding to each channel in the image and their overall expression levels corresponding sequentially to the 16 channels from top to bottom in image (B). The standard deviation represents different positions within each channel (n = 5). (**D**) Representative images for each marker in the DLBCL2 sample and the corresponding diagnosis provided by pathologists based on these images. '+' denotes positivity in tumor cells; '−' denotes negativity in tumor cells.

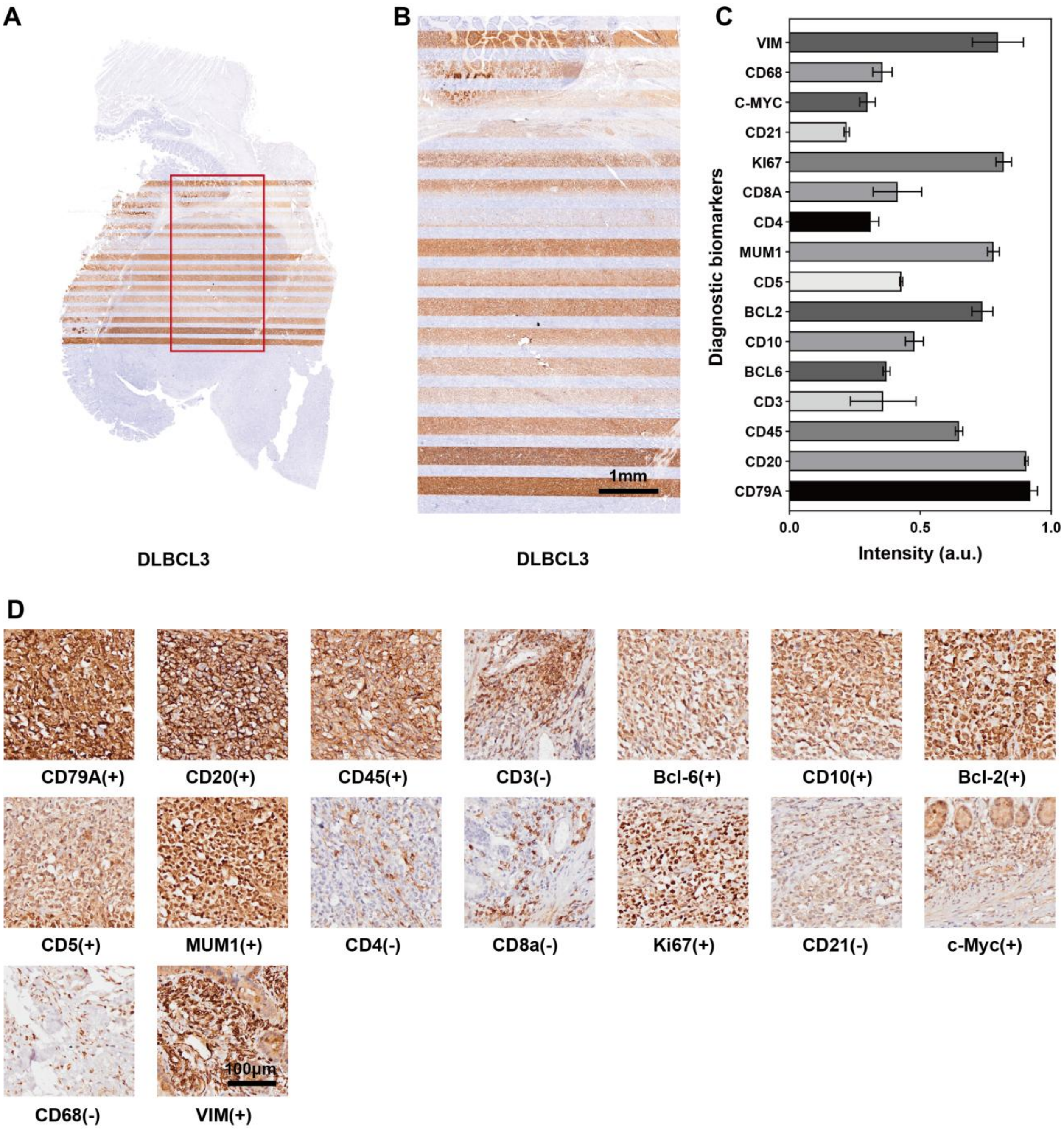


**Fig. S10. Single slide diagnostic markers expression for DLBCL3.** (**A**) DLBCL3 sample after Dumi-based multi-channel mIHC staining. (**B**) Magnified view of the mIHC image. (**C**) Markers corresponding to each channel in the image and their overall expression levels corresponding sequentially to the 16 channels from top to bottom in image (B). The standard deviation represents different positions within each channel (n = 5). (**D**) Representative images for each marker in the DLBCL3 sample and the corresponding diagnosis provided by pathologists based on these images. '+' denotes positivity in tumor cells; '−' denotes negativity in tumor cells.

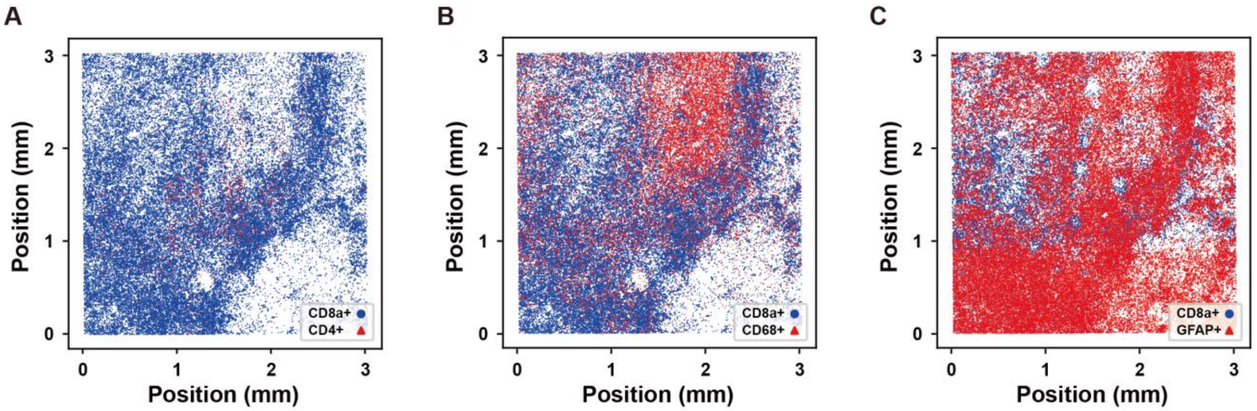


**Fig. S11. Spatial distributions of CD8a+ and other cell classifications in PCNS-DLBCL.** (**A**) Spatial distribution of CD8a+ and CD4+ cells within the region shown in Fig. 6A. (**B**) Spatial distribution of CD8a+ and CD68+ cells within the region shown in Fig. 6A. (**C**) Spatial distribution of CD8a+ and GFAP+ cells within the region shown in Fig. 6A.

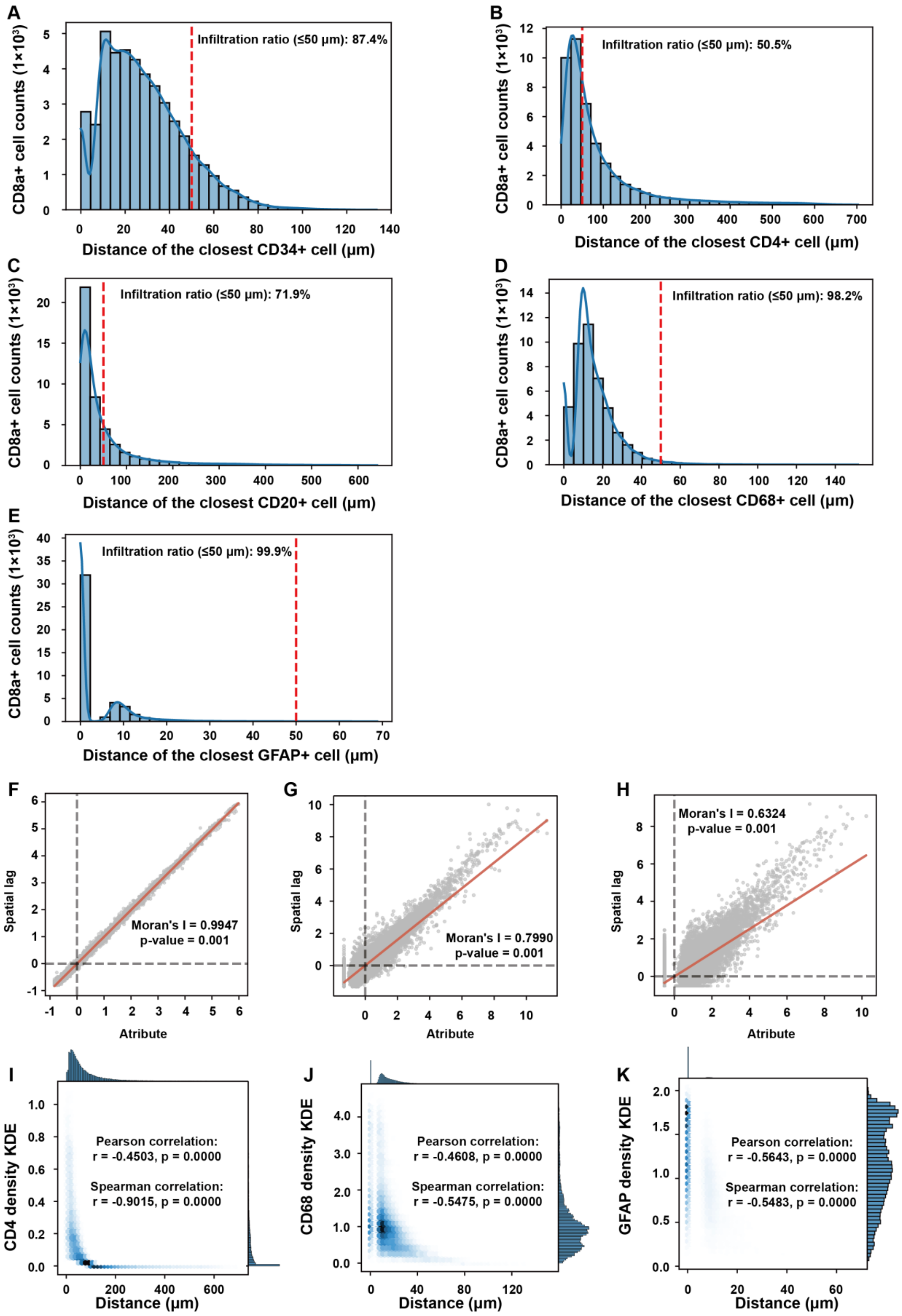
A
Infiltration ratio (≤50 μm): 87.4%
CD8a+ cell counts (1×10³)
Distance of the closest CD34+ cell (μm)
B
Infiltration ratio (≤50 μm): 50.5%
CD8a+ cell counts (1×10³)
Distance of the closest CD4+ cell (μm)
C
Infiltration ratio (≤50 μm): 71.9%
CD8a+ cell counts (1×10³)
Distance of the closest CD20+ cell (μm)
D
Infiltration ratio (≤50 μm): 98.2%
CD8a+ cell counts (1×10³)
Distance of the closest CD68+ cell (μm)
E
Infiltration ratio (≤50 μm): 99.9%
CD8a+ cell counts (1×10³)
Distance of the closest GFAP+ cell (μm)
F
Moran's I = 0.9947
p-value = 0.001
Spatial lag
Atribute
G
Moran's I = 0.7990
p-value = 0.001
Spatial lag
Atribute
H
Moran's I = 0.6324
p-value = 0.001
Spatial lag
Atribute
I
Pearson correlation:
r = -0.4503, p = 0.0000
Spearman correlation:
r = -0.9015, p = 0.0000
CD4 density KDE
Distance (μm)
J
Pearson correlation:
r = -0.4608, p = 0.0000
Spearman correlation:
r = -0.5475, p = 0.0000
CD68 density KDE
Distance (μm)
K
Pearson correlation:
r = -0.5643, p = 0.0000
Spearman correlation:
r = -0.5483, p = 0.0000
GFAP density KDE
Distance (μm)

**Fig. S12. Analysis of relative spatial distributions between CD8a+ cells and other populations in PCNS-DLBCL.** (**A–E**) Histograms of nearest-neighbor distances from CD8a$^+$ cells to CD34+, CD4+, CD20+, CD68+, and GFAP+ cells; red dashed line at 50 μm denotes the threshold for infiltration index. (**F–H**) Spatial autocorrelation (Moran's I) analysis of nearest-neighbor distances from CD8a+ cells to each cell type; red solid line is the global regression line with slope equal to Moran's I. (**I–K**) KDE joint distribution plots of nearest neighbor distances from CD8a+ to CD4+, CD68+, and GFAP+ cells versus local densities of those cell types.

**Table S1. Time comparion between Dumi-based immunostaining and conventional immunostaining (only list the key steps that are different).**

| reagent | Conventional IHC | Dumi-based IHC | Conventional TSA-based mIF (one round) | TSA-based mIF processed by Dumi (one round) |
|---|---|---|---|---|
| Hydrogenperoxide blocking | 25 min | 5 min | 25 min | 5 min |
| Washing buffer | 5 min | 30 s | 5 min | 30 s |
| Blocking solution | 30 min | 8 min | 30 min | 8 min |
| Primary antibody treatment | 1 h–12 h | 10 min | 1 h–12 h | 10 min |
| Washing buffer | 5 min | 30 s | 5 min | 30 s |
| Secondary antibody treatment | 30 min−1 h | 8 min | 30 min−1 h | 8 min |
| Washing buffer | 5 min | 30 s | 5 min | 30 s |
| DAB or TSA dye | 10 min | 1 min | 10 min | 1 min |
| DI water | 5 min | 30 s | 5 min | 30 s |
| Total consumed time | 175 min (at least) | 34 min | 175 min(at least) | 34 min |

**Table S2. CD20 two-way ANOVA analysis result.**

| ANOVA table | % of total variation | SS | DF | MS | F (DFn, DFd) | P value |
|---|---|---|---|---|---|---|
| Interaction | 5.010 | 0.1528 | 10 | 0.01011 | $F (10, 342) = 63.70$ | $p<0.0001$ |
| Time Factor | 53.41 | 0.8474 | 5 | 0.2155 | $F (5, 342) = 1358$ | $p <0.0001$ |
| Flow Factor | 38.89 | 1.759 | 2 | 0.3922 | $F (2, 342) = 2472$ | $p <0.0001$ |
| Residual | | 0.04290 | 342 | 0.0001587 | | |

Additional Tukey's HSD analyses showed that, at the same flow rate, only the comparison between 25 min and 30 min in the 4 μL min$^{-1}$ group failed to reach statistical significance ($p > 0.05$), whereas at the same incubation time, only the comparison between 2 μL min$^{-1}$ and 4 μL min$^{-1}$ in the 5 min group was not significant ($p > 0.05$).

**Table S3. Ki67 two-way ANOVA analysis result.**

| ANOVA table | % of total variation | SS | DF | MS | F (DFn, DFd) | P value |
|---|---|---|---|---|---|---|
| Interaction | 5.451 | 0.1528 | 10 | 0.01528 | $F (10, 342) = 121.8$ | $p < 0.0001$ |
| Time Factor | 30.24 | 0.8474 | 5 | 0.1695 | $F (5, 342) = 1351$ | $p < 0.0001$ |
| Flow Factor | 62.78 | 1.759 | 2 | 0.8797 | $F (2, 342) = 7012$ | $p < 0.0001$ |
| Residual | | 0.04290 | 342 | 0.0001254 | | |

Additional Tukey's HSD analyses showed that, within the 2 μL min$^{-1}$ condition, the 10 min vs. 15 min, 20 min vs. 25 min, and 25 min vs. 30 min comparisons; within the 4 μL min$^{-1}$ condition, the 15 min vs. 20 min, 15 min vs. 25 min, 15 min vs. 30 min, 20 min vs. 25 min, 20 min vs. 30 min, and 25 min vs. 30 min comparisons; and within the 6 μL min$^{-1}$ condition, only the 20 min vs. 25 min comparison, did not achieve statistical significance ($p > 0.05$).

**Table S4. Primary Antibody Panel.**

| Marker | Species | Company | Catalog Number | Origin |
|---|---|---|---|---|
| CD20 | Mouse mAb | Servicebio | GB14030-50 | China |
| Ki67 | Rabbit pAb | Servicebio | GB111499-100 | China |
| CD8a | Mouse mAb | Servicebio | GB12068-100 | China |
| CD68 | Rabbit pAb | Servicebio | GB113150 | China |
| CD4 | Rabbit mAb | Boster | BM4263 | China |
| CD34 | Rabbit mAb | Boster | BM4082 | China |
| CD79A | Rabbit pAb | Boster | PB9168 | China |
| CD45 | Mouse mAb | Servicebio | GB14038-50 | China |
| CD3 | Rabbit pAb | Servicebio | GB11014-50 | China |
| CD5 | Rabbit pAb | Boster | A00480-2 | China |
| CD10 | Rabbit pAb | Servicebio | GB114689-50 | China |
| Bcl-2 | Rabbit mAb | Servicebio | GB154380-50 | China |
| Bcl-6 | Rabbit mAb | Boster | BM4070 | China |
| c-Myc | Rabbit mAb | Boster | BM4042 | China |
| MUM1 | Rabbit pAb | Boster | PB9222 | China |
| VIM | Rabbit pAb | Servicebio | GB11192-50 | China |
| CD21 | Mouse mAb | Servicebio | GB14031-50 | China |
| GFAP | Mouse mAb | Servicebio | GB12100-50 | China |
